# Comparative analysis of centromeres of oat (*Avena sativa*) and its tetraploid and diploid relatives reveals rapid evolution of centromere composition and architecture

**DOI:** 10.1101/2025.04.08.647780

**Authors:** Carlotta Marie Wehrkamp, Matthias Heuberger, Thomas Wicker

## Abstract

**Background:** Recent advances in assemblies of nearly gap-free, high-quality genomes have enabled detailed analysis of centromeres in large and highly repetitive crop genomes. Here, we analyse the hexaploid *Avena sativa* genome and its tetraploid (*A. insularis*) and diploid relatives (*A. longiglumis*, *A. atlantica* and *A. eriantha*).

**Results:** *Avena* centromeres are largely composed of retrotransposons belonging to three families, *RLG_Ava*, *RLG_Cereba* and *RLG_Beth*. Analysis of retrotransposon populations revealed striking differences in the centromere architecture between the A, C and D genome lineages. We identified distinct profiles of transposable element bursts for these lineages which include the emergence and disappearance of retrotransposon families and subfamilies. We identified multiple centromere shifts which occurred in the C genome lineage within the past ∼ 4 myrs and retraced species divergences and polyploidization events in the *Avena* genus. Although the studied species are closely related, our data show that their centromeres have rapidly evolved different and distinct centromere architectures, for example through the spread of novel satellite repeats or activity bursts of different retrotransposon families and subfamilies. Additionally, we found that *RLG_Ava* and *RLG_Cereba* retrotransposons have been coexisting while simultaneously competing for the centromeric “niche” since the emergence of the *Poaceae* (grasses).

**Conclusions:** Our comparative analyses provided detailed insight into centromere evolution across the *Avena* genus and revealed that composition and architecture of centromeres can vary greatly even between closely related species and different ploidy levels. Our findings emphasize the need for extended analyses of large genome species to improve our understanding of centromere evolution.

## Background

Centromeres are essential in the cell division of eukaryotes. They are the assembly site of the kinetochore to which microtubules bind during mitosis and meiosis and thus mediate chromosome segregation. In most species centromeres are defined epigenetically, rather than through a specific sequence of DNA, by the presence of the centromere specific CENH3 histone variant, which substitute two canonical H3 histones [1,2]. There are two main types of centromeres: Holocentromeres, in which the centromere stretches along the whole chromosome or where the spindle fibers attach the multiple CENH3 clusters (cluster-like holocentromere) and monocentromeres, in which a single CENH3 containing region defines the functional centromere [3].

The *Poaceae* (grasses) family comprises all major cereal crops, for example wheat, barley, oats, maize, rye and rice. The most common centromere architecture found within the grass family are “regional” or monocentromeres, i.e. centromeres that comprise a defined, small part of chromosomes and are usually a few Mb in size [4,5]. First evidence of centromere specific repetitive sequences in cereals was brought up in 1996 by Jiang et al. [6]. They demonstrated the presence of pSau3A9 (isolated from *Sorghum bicolor*) in the centromeric regions of various grass chromosomes including oat. In the same year Aragón-Alcaide et al. showed that the CCS1 (cereal centromeric sequence, isolated from *Brachypodium*) family occurred in the centromeres of the Triticeae, maize and rice [7]. Both sequences were later shown to be part of a centromere specific retrotransposon belonging to the *Gypsy* superfamily [8,9]. They are conserved in the centromeres of grasses [8] and have repeatedly been shown to localize in the functional centromere by fluorescence in-situ hybridization assays (FISH) and chromatin immunoprecipitation (ChiP) (e.g. rice and maize) [10–12]. Most centromere-specific retrotransposons belong to an ancient clade of *Gypsy* elements [13]. These are called CRM in maize, CRR in rice and *Cereba* in barley [9,11,12].

In recent years, the tribe of the Triticeae, which includes some of the world’s most important crops such as wheat, barley and rye, has become a model for the study of large and highly repetitive genomes. The dominant centromere-specific transposable element (TE) family in Triticeae is *RLG_Cereba* [14–17]. Curiously, centromeres of Triticeae also contain a second, less abundant transposable element (TE) family called *RLG_Quinta*. *RLG_Quinta* is a non-autonomous retrotransposon, i.e. a TE family which does not encode canonical retrotransposon proteins and which likely evolved from *RLG_Cereba* elements through loss of most of its coding sequences [18].

It has long been speculated that centromere specificity of centromere-specific TEs is due to a unique protein domain that is fused to the C-terminus of the integrase enzyme (called CR-domain) [13,19,20]. Recently, Heuberger et al. [18] proposed that the centromere specificity of *RLG_Cereba* might be rooted in a conserved RARA[RK]-motive in the CR-domain directly interacting with the CENH3 histone variant. Additionally, the study proposed that the long terminal repeats (LTRs) of *RLG_Cereba* and its non-autonomous partner, *RLG_Quinta*, are involved in the phasing/placement of CENH3 containing nucleosomes.

Only a few *Triticeae* genomes have completely sequenced, gap-free centromeres (e.g. *T. monococcum*, [16]). While centromeres of rye and several wheat genomes are partially sequenced [15,21], the centromeres of barley remain largely unresolved due to the presence of highly repetitive tandem satellite repeats [22].

Many questions therefore remain unanswered about centromeres in the large-genome cereals. For example, hexaploid wheat contains an additional centromere-specific retrotransposon family called *RLG_Abia* [23]. However, these elements seem to have been silent during recent evolutionary times, as mostly fragments are found. Considering the rapid turnover of repetitive sequences in grasses (amplification of TEs and their continuous removal through deletions; [24]), this indicates that *RLG_Abia* elements became silent only relatively recently and suggests that different TE families may populate cereal centromeres at different times.

Recently, a highly contiguous genome of the hexaploid oat *Avena sativa* (ACD) was made available (Avena sativa – OT3098 v2, PepsiCo, https://wheat.pw.usda.gov/jb?data=/ggds/oat-ot3098v2-pepsico). With a size of over 10 Gb, it is among the largest genomes sequenced so far. This was also the first assembly in which centromeres were assembled nearly gap free. Centromeres of the A and D subgenome were shown to consists mainly of LTR retrotransposons, while C subgenome centromeres also contained highly abundant tandem repeats [25]. Oat (*A. sativa*) is a close cereal relative to the Triticeae tribe which contains wheat, barley and rye. The Aveneae and Triticeae lineages diverged approximately 28 million years ago [26] while the Triticeae themselves radiated approximately 8-12 million years ago [26–28]. The genus *Avena* includes diploid, tetraploid and hexaploid species. *A. sativa* is an agronomically important allohexaploid crop (genome size = 10.78 Gb) comprising three large and highly repetitive A, C and D subgenomes.

The phylogeny of hexaploid oat has long been a topic of discussion. Different *Avena* species have been brought up as potential donors of the genomes found in hexaploid oats and especially the origin of the D genome has been the focus of recent studies. The A and D genome are thought to be more closely related to each other than to the C genome [29,30]. In addition, it has been suggested that the D genome of hexaploid oats is derived from the AC tetraploids. And further, based on genome wide analysis and phylogenetic studies, it was proposed that these tetraploids carry a D genome instead of an A genome [31–34]. The tetraploid *Avena insularis* (CD, formerly AC) is presumed to be the progenitor of hexaploid oats with evidence being provided for example by chromosome morphology similarity [35,36], through genetic diversity studies [37], high-density genotyping-by-sequencing markers [32] and recently by cytogenetic analysis using FISH [34].

On the other hand, a phylogenetic study from 2018 based on the *Pkg1* (nuclear plastid 3-phosphoglycerate kinase) gene, suggest that the tetraploids *Avena murphyi* or *Avena marrocana* might be a closer relative to the C and D genomes found in hexaploid oats [38], however the consensus seems to emerge that *A. insularis* is the probable CD genome donor. A high-quality reference genome for *A. insularis* was recently presented by Kamal et al. [39].

The origin of the A genome of hexaploid oats has been studied extensively, however no particular *Avena* species has been widely agreed on as a putative donor yet. Some studies analyzed the diploid *Avena longiglumis* (A_l_) [26,39,40] as a possible progenitor of the A genome in hexaploid oat. Others highlighted that *Avena atlantica* (A_s_) is closely related to the A genome in hexaploid oat and discussed both of the above mentioned diploids as putative diploid ancestors [26,30,38,40]. Reference genomes were made available recently for *A. longiglumis* [39] and *A. atlantica* [41].

Using the above mentioned genomes from the *Avena* genus, the goal of this study was to characterize the centromeric architecture in this genus, with particular focus on centromeric TEs. We found that *Avena* centromeres are dominated by two putatively autonomous retrotransposon families, *RLG_Cereba* and *RLG_Ava*, which seem to compete for the centromeric „niche“ in different *Avena* species. We use these two families to investigate the evolutionary history of multiple *Avena* genomes, with a focus on previously proposed progenitors of hexaploid oat. Finally, we explored the evolutionary origins of *RLG*_*Ava* and *RLG_Cereba* elements in the *Poaceae* and demonstrated their coexistence for millions of years.

## Results

### Two autonomous and one non-autonomous retrotransposon family are abundant in the centromeres of *A. sativa*

A recent study determined positions of functional centromeres in the *A. sativa* cv. Sang genome assembly using CENH3 ChIP-seq data [42]. Since we did not have ChIP-seq data available, positions of functional centromeres in the *A. sativa* cv. OT3098 genome were predicted by projecting the centromere positions of the *A. sativa* cv. Sang genome on to the OT3098 assembly (Example in Fig. S1, Tab. S1). Because we expected centromeric TEs similar to those previously described in grasses [9,12,16,43] we searched for retrotransposons encoding an integrase with a CR-domain, which has recently been described to be important for the centromere specificity of these elements [13,18–20].

We identified two main centromere-specific families, one homologous to the known *RLG_Cereba* (here named *RLG_Cereba*) and one similar to the previously described *RLG_Abia* elements (here named *RLG_Ava*, Fig. 1a and b). Both are putatively autonomous families, meaning they encode all proteins necessary for transposition. In addition, both families feature a CR domain with a RARA[RK]-motive located in the beginning of the 3’ LTR (Fig. 1a and b, see below).

**Figure 1.**
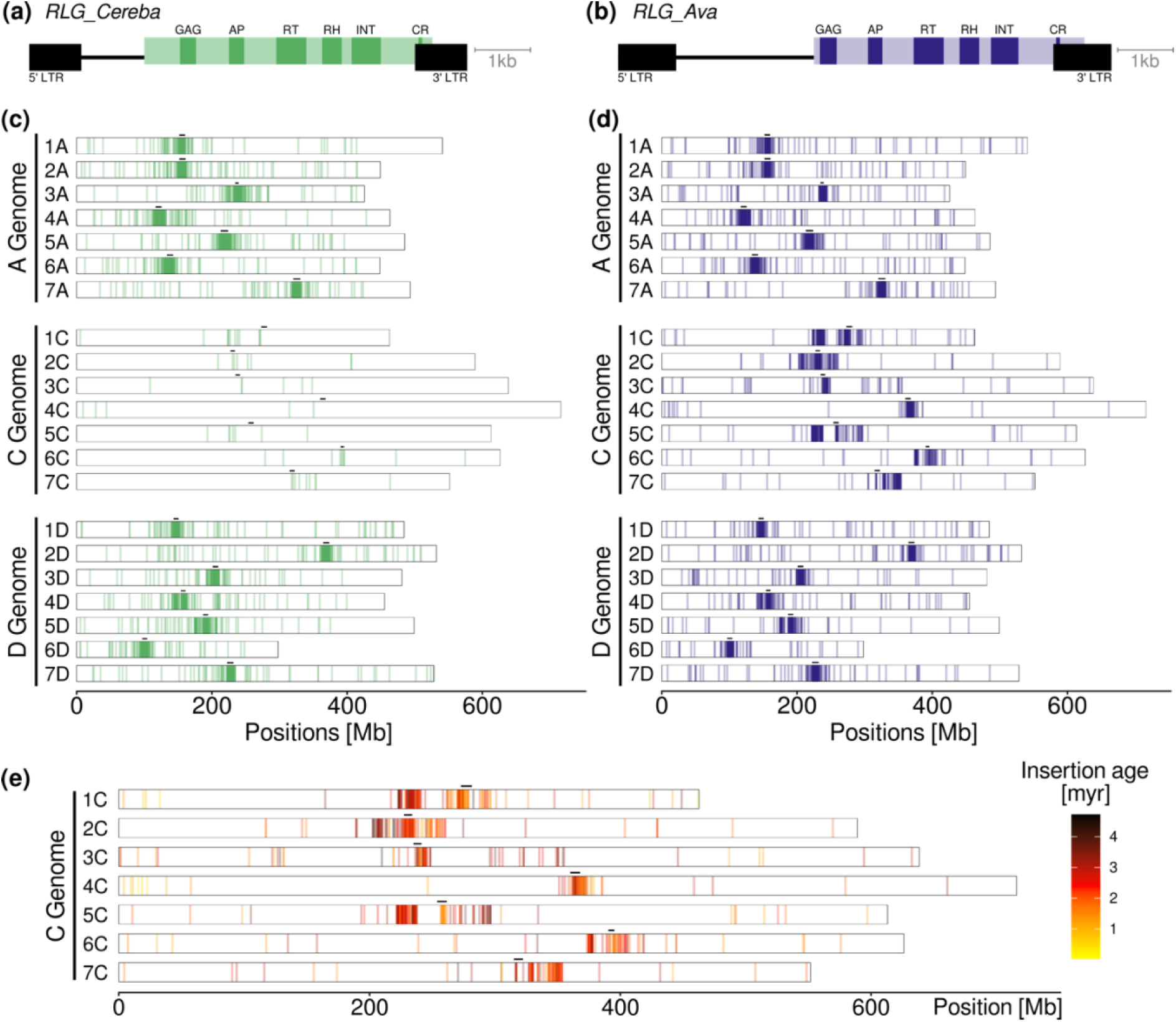
Distribution of centromere specific retrotransposons in *A. sativa* OT3098. Sequence organization of *RLG_Cereba* **(a)** and *RLG_Ava* (from C subgenome) elements **(b)** AP: aspartate protease, RT: reverse transcriptase, RH: RNase H, INT: integrase, CR: Chromodomain. **(c)** Chromosomal distribution of full-length *RLG_Cereba* retrotransposons. Predicted positions of functional centromeres are indicated by black bars. Note that the C subgenome contains only very low numbers of *RLG_Cereba* elements. **(d)** Chromosomal distribution of *RLG_Ava* retrotransposons. **(e)** Insertion age and distribution of *RLG_Ava* elements in subgenome C.

When comparing consensus sequences of *RLG_Cereba* and *RLG_Ava*, we found the conservation to be too low to reliably align them on DNA level. Their predicted proteins show a similarity of 65.2% to 68.5%, indicating that these families represent distinct and ancient evolutionary lineages. Additionally, we identified multiple variants of *RLG_Ava* elements which differ strongly in sequences of short tandem repeat clusters in their LTRs (example in Fig. S2). Interestingly, we identified copies of the *RLG_Cereba* family almost exclusively in the A and the D subgenomes (Fig. 1c). We found 1209 and 1079 *RLG_Cereba* full-length copies in the A and D subgenome, respectively, while in the C subgenome we identified only 57 copies. In contrast, we found high numbers of *RLG_Ava* elements in all oat subgenomes, with 1,468 copies in A, 745 in C and 1,076 copies in the D subgenome (Fig. 1d; Fig. S3).

For all full-length copies, we estimated their insertion age based on divergence of their LTRs [44]. Average insertion age of *RLG_Cereba* elements was estimated to be approximately 0.89 myrs on the A genome and 1.04 myrs in the D genome, while the 57 copies on the C genomes were on average 2.17 myrs old. Results were similar for *RLG_Ava* elements: while the A and D subgenome comprised elements with average ages of around 0.81 and 1.02 myrs, respectively, copies from the C subgenome were much older, with an average insertion age of 2.29 myrs (Fig. 1e).

In addition to *RLG_Cereba* and *RLG_Ava*, we identified another retrotransposon family (*RLG_Beth*), of which we found 1,180, 506 and 1,000 full-length copies on the A, C and D subgenome, respectively. *RLG_Beth* copies also localize in the centromeric region of the chromosomes (Fig. S4). *RLG_Beth* encodes a large protein of unknown function, but does not encode the proteins required for transposition. We propose that *RLG_Beth* is a non-autonomous partner family to *RLG_Ava*, and relies on the autonomous partner for its replication. Indeed, *RLG_Beth* and *RLG_Ava* show sequence conservation in their LTRs and primer binding site, previously described characteristics of autonomous/non-autonomous pairs [17,45]. One example is shown in Fig. S5 for TEs from the subgenome D. Furthermore, overall copy numbers and insertion ages of *RLG_Beth* elements are similar to those of *RLG_Ava* elements: copies in the C subgenome were estimated to be ∼2.26 myrs old on average, while in the A and D subgenome the copies showed a lower insertion age of ∼1.22 myrs and ∼1.23 myrs respectively.

### Centromere-specific TEs indicate the positions of functional centromeres

We analyzed the positions of the most recent insertions of *RLG_Cereba*, because it was shown in wheat that the youngest *RLG_Cereba* copies were found inside or close to the functional centromere [16]. In oat, we found that most of the recently inserted *RLG_Cereba* elements, i.e. elements younger than 1 myrs, are highly enriched in the predicted functional centromeres on subgenomes A and D (Fig. S6). Likewise, *RLG_Ava* retrotransposons with insertion ages younger than 1 myr were consistently enriched in predicted functional centromeres. In fact, insertion sites of the youngest copies for both *RLG_Cerea* and *RLG_Ava* perfectly coincided with centromere position inferred from CENH3 ChIP-seq data [42] in all chromosomes of the A an D subgenomes (Fig. S6 and S7). We thus concluded that young insertion of both *RLG_Cerea* and *RLG_Ava* are highly reliable markers for functional centromeres.

However, the 745 *RLG_Ava* copies from the C subgenome were much older, with an average insertion age of ∼2.29 myrs, while *RLG_Cereba* retrotransposons are virtually absent from the C genome. We therefore wanted to study whether the older elements also mark functional centromere positions. With the exception of elements localised in the previously described translocations on 1C and 4C (e.g., [29,39,46–48]), we found the youngest elements (in this case younger than 2 myrs) to accumulate within or at the sites of the predicted centromeres in chromosomes 1C, 4C, 5C and 6C (Fig. S7). In chromosome 7C, the aggregation of the young elements did not colocalize with the predicted centromere position. On chromosome 2C, we found no distinct accumulation site but rather a broader distribution of young *RLG_Ava* elements in the region of the predicted centromere. Similarly, on chromosome 3C we identified only few elements and therefore also were not able to draw conclusions on the position of the functional centromere (Fig. S7).

In summary, centromeric TEs in the A and D subgenomes are clearly younger than those in the C subgenome, many of them having an insertion age of 0 (i.e. identical LTRs). This indicates that A and D subgenome centromeres are still actively being reshuffled, similar to those in Einkorn wheat [16]. Consequently, insertion of the youngest TE copies coincided well with the inferred positions of the centromeres. In contrast, centromeric TEs in the C subgenome seem to have been largely “silent” for a long time, suggesting that different dynamics are influencing positions of functional centromeres in the C subgenome.

### *RLG_Ava* subfamilies reveal centromere shifts in the *A. sativa* C subgenome

The observations that *RLG_Ava* retrotransposons are much older in the C subgenome and that predicted positions of functional centromeres differ in some cases from accumulations of youngest *RLG_Ava* elements inspired us to study the evolution of *RLG_Ava* elements in more detail. Using a TE population analysis pipeline [45] we identified eight subfamilies of *RLG_Ava* among the 745 full-length copies in the C subgenome. Four subfamilies which comprised more than 50 elements were included in our further analysis. All subfamilies were found enriched in the centromeric and pericentromeric regions of the C subgenome and mostly clustered by their ages per chromosome (Fig. 2a and b).

**Figure 2.**
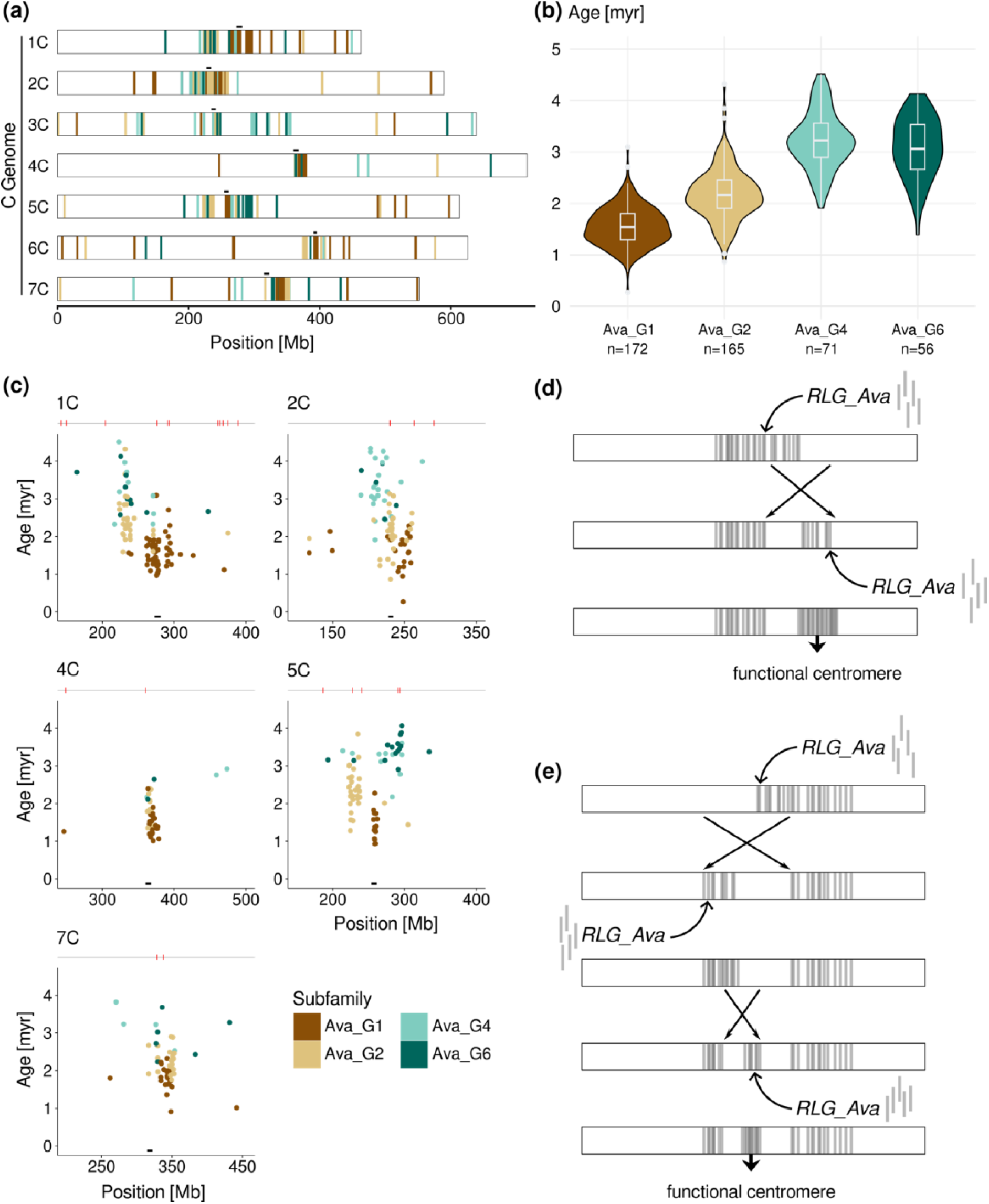
*RLG_Ava* subfamilies reveal centromere shift in *A. sativa* OT3098 C subgenome. The distribution of *RLG_Ava* elements **(a)** and the respective age distribution of the different subfamilies **(b)**. Age distributions of four diagnostic *RLG_Ava* subfamilies. **(c)** Insertion ages of *RLG_Ava* elements across the position within the centromeric region of *A. sativa* OT3098 for 5 chromosomes. Each dot represents one retrotransposon copy. The red marks on the gray track indicate the positions of gaps in the OT3098 assembly. The black bar indicates the projected centromere position. **(d)** Evolutionary model for an inversion leading to a centromere shift in chromosome 1C. A large inversion of ∼30 Mb moves part of the centromere to the right where a new functional centromere is established. **(e)** Evolutionary model for the centromere shift due to two successive inversions in chromosome 5C.

Subfamilies G4 and G6 were most active ∼3 mya, while G1 and G2 had their peak activity ∼2.2 and ∼1.5 mya, respectively. Interestingly, we observed that the focus of the insertion sites of *RLG_Ava* subfamilies changed over the course of the last ∼4 myrs in some C subgenome chromosomes. While the positions of the functional centromeres appear to be steady in chromosomes 4C and 7C (Fig. 2c), on chromosome 2C, there is a gradual shift of this preferential site towards the middle of the chromosome during the past ∼4 myrs (Fig. 2c). A more dramatic centromere shift was found in chromosome 1C, which is likely the result of an inversion transferring part of the formerly active centromere more towards one end of the chromosome where, in the following, the now functional centromere was established (Fig. 2d; Fig. S8). Additionally, it appears that there have been two shifts of the centromere on chromosome 5C within the last ∼4 myrs. The first one around 3 to 3.5 mya brought the centromeric region from a position around 290-296 Mb to roughly 222-238 Mb. The second one took place approximately 1.5 to 2 mya and brought the centromere to the site we currently predict the functional centromere at and which simultaneously harbours the youngest group of *RLG_Ava* elements (Fig. 2a, c and e; Fig. S8). These observed centromere shifts in 1C and 5C can not be explained through mis-assemblies as the two centromeres are nearly free of sequence gaps (Fig. 2c; Fig. S8 and S9). In addition, we found these centromere shifts also when analyzing the *RLG_Beth* subfamilies on the C subgenome (Fig. S10).

The centromere on chromosome 7C also remained steady, however, like for chromosome 2C, the predicted centromere position diverges from the localisation of the youngest *RLG_Ava* copies (Fig. 2c), suggesting other factors than retrotransposon sequences to determine the position of centromeres. For the centromeres of chromosome 3C and 6C, we could not identify enough full-length copies of *RLG_Ava* to allow further conclusions on their dynamics in the last ∼4 million years, although in the case of the 3C chromosome it can be stated that the youngest elements present within the overall centromeric region co-localize with the predicted function centromere position (Fig. 2a).

### Analysis of *RLG_Ava* and *RLG_Cereba* sheds light on *Avena* subgenome phylogeny

To study TE and centromere dynamics in the *Avena* genus, we expanded our analysis and included genome assemblies from two A-genome diploids, *A. longiglumis* and *A. atlantica* which carry the A_l_ and A_s_ genomes, respectively. Additionally, we include the genome of *A. insularis* (the presumed CD tetraploid progenitor of hexaploid oats) and *A. eriantha*, a diploid C genome relative carrying the C_p_ genome. We once more utilized the TE population analysis pipeline [45] and analyzed the *RLG_Ava* and *RLG_Cereba* families separately.

For *RLG_Ava*, the initial analysis revealed the identified 7,841 full-length copies to cluster into three distinct groups, reflecting the three subgenomes (*RLG_Ava_G1* through *G3*; Fig. S11). The three groups we divided up for a more detailed analysis which led to the identification of six subfamilies (Fig. 3). For *RLG_Cereba*, from a total of 5,486 full-length elements, we selected a subset which comprises the younger copies, as the subfamilies identified from using this subset were more diagnostic for the different genomes (Fig. S12). From this subset of 3,086 full-length copies, we also defined six subfamilies (Fig. 3).

**Figure 3.**
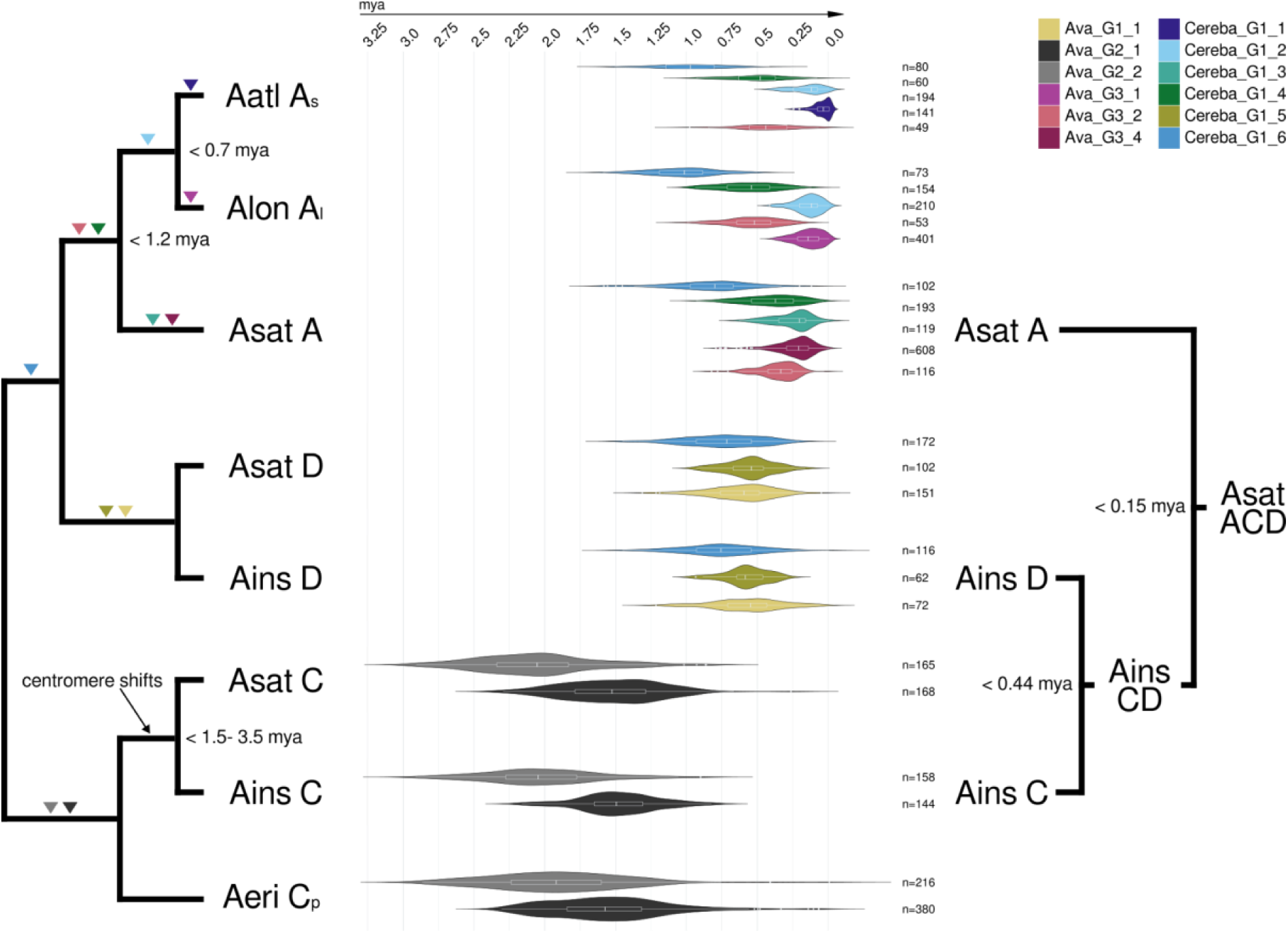
Graphic of the genetic relationship of different *Avena* species. The graphic displays the species *A. sativa* (ACD), *A. insularis* (CD), *A. longiglumis* (A_l_), *A. atlantica* (A_s_) and *A. eriantha* (C_p_). The triangles indicate the presence of a subfamily. The violin plots highlight the time of activity bursts of the respective subfamily in the different genomes. The names of the subfamilies were shortened.

Particularly informative was subfamily *RLG_Cereba_G1_6*, which was found in all A and D genomes, indicating that it was present already in a common ancestor of the A and D genomes. The A/D genome specific presence of the subfamily is in line with previous findings that that the A and D are more closely related to each other than to the C genome genome [29,30]. Our analysis indicates that *RLG_Cereba_G1_6* had independent activity bursts in the A and D genome lineages between ∼0.5 and ∼1.2 myr ago (Fig. 3). Here, we defined the duration of a TE activity burst as the time span that covers the inter quartile range (IQR, 25 ^th^ to 75^th^ percentile) of the insertion ages of a given subfamily. We know that the activity bursts in the A and D genome lineages must have been independent, as they overlap in time with activity bursts of subfamilies *RLG_Cereba_G1_5* and *RLG_Ava_G1_1* which are only found in the D genomes (Fig. 3). Therefore, the A and D genome lineages must have diverged before the *RLG_Cereba_G1_6* activity burst, at least ∼1.2 myrs ago. For the D genomes of *A. sativa* and *A. insularis*, TE activity profiles of the three studied subfamilies look very similar (Fig. 3). This was expected, since *A. insularis* is the proposed progenitor of the hexaploid oats.

Interestingly, TE activity profiles in the three A genomes differed strongly, and therefore provided information about their evolutionary history. For subfamily *RLG_Cereba_G1_4*, we identified independent bursts between the A genomes. In the *A. sativa* the beginning of the burst was estimated to ∼0.55 mya while in *A. atlantica* it started ∼0.64 and in *A. longiglumis* ∼0.72 mya. Again, we conclude this must have been independent bursts because they overlap in time with activity bursts of subfamilies *RLG_Ava_G3_4* and *RLG_Cereba_G1_3* (Fig. 3) which are found only in *A. sativa*. This clearly distinguishes the *A. sativa* A subgenome from *A. longiglumis* and *A. atlantica*, indicating that the *A. sativa* A subgenome diverged from the other two ∼1.2 myr ago.

In addition, we identified multiple subfamilies of *RLG_Cereba* and *RLG_Ava* that distinguish *A. longiglumis* and *A. atlantica.* The subfamily *RLG_Cereba_G1_1* which had an activity burst in the past 100,000 years is only found in *A. atlantica*, while subfamily *RLG_Ava_G3_1* is only found in *A. longiglumis.* Furthermore, subfamily *RLG_Ava_G3_2* had activity bursts at different times. While in the *A. longiglumis* the burst began ∼0.65 mya, in *A. atlantica* it started later at ∼0.56 myr (Fig. 3). Taken together, the TE activity profiles indicated that *A. longiglumis* and *A. atlantica* must have diverged at least ∼0.72 mya.

Further, we identified two large subfamilies (*RLG_Ava_G2_1* and *RLG_Ava_G2_2*) that were only present on the C genomes (Fig. 3). Consistent with results described above, the *RLG_Ava* subfamilies were active much earlier than in the A and D subgenomes and seem to be silent since at least 1 myrs (Fig. 3). We performed a more comprehensive analysis of the *RLG_Ava* subfamilies from the C genome (Fig. S13). Interestingly, we found the above described centromere shifts (see Fig. 2) in the C genome of *A. sativa* and the tetraploid *A. insularis* but not in *A. eriantha* (Fig. S13). This observation is consistent with the proposed ancestry of *A. sativa*, in which *A. insularis* is the likely donor of the CD genomes (e.g., [34]). It also indicates the centromere shifts occurred after the C genome donor separated from *A. insularis*.

The activity profiles are also informative with regard to polyploidization events: the two subfamilies *RLG_Cereba_G1_5* and *RLG_Ava_G1_1* which are specific for the D genomes had activity bursts from ∼0.65-0.47 and ∼0.71-0.44 (Fig. 3). Because we do not find these families in the C genome, tetraploidization must have happened after they went silent. Thus, we propose that the tetraploidization which gave rise to *A. insularis* happened less than ∼0.44 mya. Conversely, we did not find copies of the A genome specific subfamily *RLG_Ava_G3_4* in the C and D genome of *A. sativa*. Thus, we conclude that the hexaploidization (ACD) occurred no more than ∼0.15 mya, as the burst of this family ended around that time in the *A. sativa* A genome.

The molecular dating of polyploidization events is complemented by the LTR retrotransposon family *RLG_Aurora* which is not centromere specific. *RLG_Aurora* is the only TE family that we found to have an activity burst specifically in the C and D subgenomes (i.e. it was active after the tetraploid formed and became silent before hexaploidization, Fig. S14). Indeed, *RLG_Aurora* is found in all diploid genomes only at very low copy numbers, but has over 1,200 copies in the tetraploid. *RLG_Aurora* was most active between ∼0.41 mya and ∼0.36 mya in the C and D genome respectively. Using the IQR as a marker we estimated the burst to have happened from around 0.46 to ∼0.26 mya, which fits into the period between tetra-and hexaploidization estimated above, as this time of the burst is located within the estimated time window between the tetraploidization (CD) and hexaploidization (ACD). *RLG_Aurora* is thus one of the few documented examples of a TE family being activated after polyploidization.

Taken together, the comprehensive analysis of two centromere specific retrotransposon families strongly supports hypothesis that *A. insularis* is the CD genome donor of hexaploid oats [34,39]. For the *A. sativa* A subgenome, we conclude that it is basal to *A. longiglumis* and *A. atlantica*, as the latter two share more similarities in their TE activity profiles and are therefore closer related to one another than to the *A. sativa* A subgenome.

### C subgenome centromeres have a large fraction of tandem repeats

To study the overall composition of *Avena* centromeres we annotated them with EDTA for TEs [49] and TRF for tandem repeats [50]. The A and D subgenomes have low tandem repeat contents of 3.6% and 2.9%, respectively. In contrast, the C subgenome has a more than 5-fold higher tandem repeat content (16.9%; Fig. S15). A majority of the tandem repeat arrays found in the C genome centromeres consist of a repeat with a period size of 46-51. Likely, this corresponds to a previously described tandem repeat [25], we however did not observe this or any other tandem repeats we identified to be specific to centromeres. All three subgenomes have a high TE content (A=92.6%, C=75.09%, D=89.9%). Most TEs are of the *Gypsy* superfamily (A=80.6%, C=50.4%, D=76.2%), of which *RLG_Ava* and *RLG_Cereba* are the most abundant. Together, tandem repeats and TEs account for 96.2% (A), 91.99% (C), and 92.8% (D) of the centromeres in *A. sativa,* meaning that the centromeric DNA is highly repetitive (Fig. S15). In summary, these analyses show that the A and D subgenomes have almost exclusively TE based centromere architectures, whereas the C subgenome differs in that it contains a considerable fraction of tandem repeats that are part of the functional centromere.

### *RLG_Ava* and *RLG_Cereba* lineages are conserved across the *Poaceae*

To further investigate the relationship between the two centromere specific family *RLG_Cereba* and *RLG_Ava*, we identified homologs in various grasses as well as more instantly related species such as ginger, pineapple, pine, the two arabidopsis species *Arabidopsis thaliana* and *Arabidopsis lyrata*. We then constructed a phylogenetic tree from the predicted proteins of these TE families using MrBayes [51]. We found the homologs of *RLG_Ava* and *RLG_Cereba* to form two distinct clades within the phylogenetic tree. For each of the investigated grass genomes we found a homolog for both families, indicating that the two TE lineages must have been present in the ancestor of all grasses (*Poaceae*) before the divergence of *Streptochaeta* ∼100 mya [52] (Fig. 4). This indicates that the two families have been coexisting while simultaneously competing for the same genomic niche. Interestingly, the *RLG_Ava* elements of the different *Avena* species cluster in an AD and the C genome clade while the *RLG_Cereba* elements from the different (sub-)genomes do not follow this pattern and are more similar to one another (Fig. 4).

**Figure 4.**
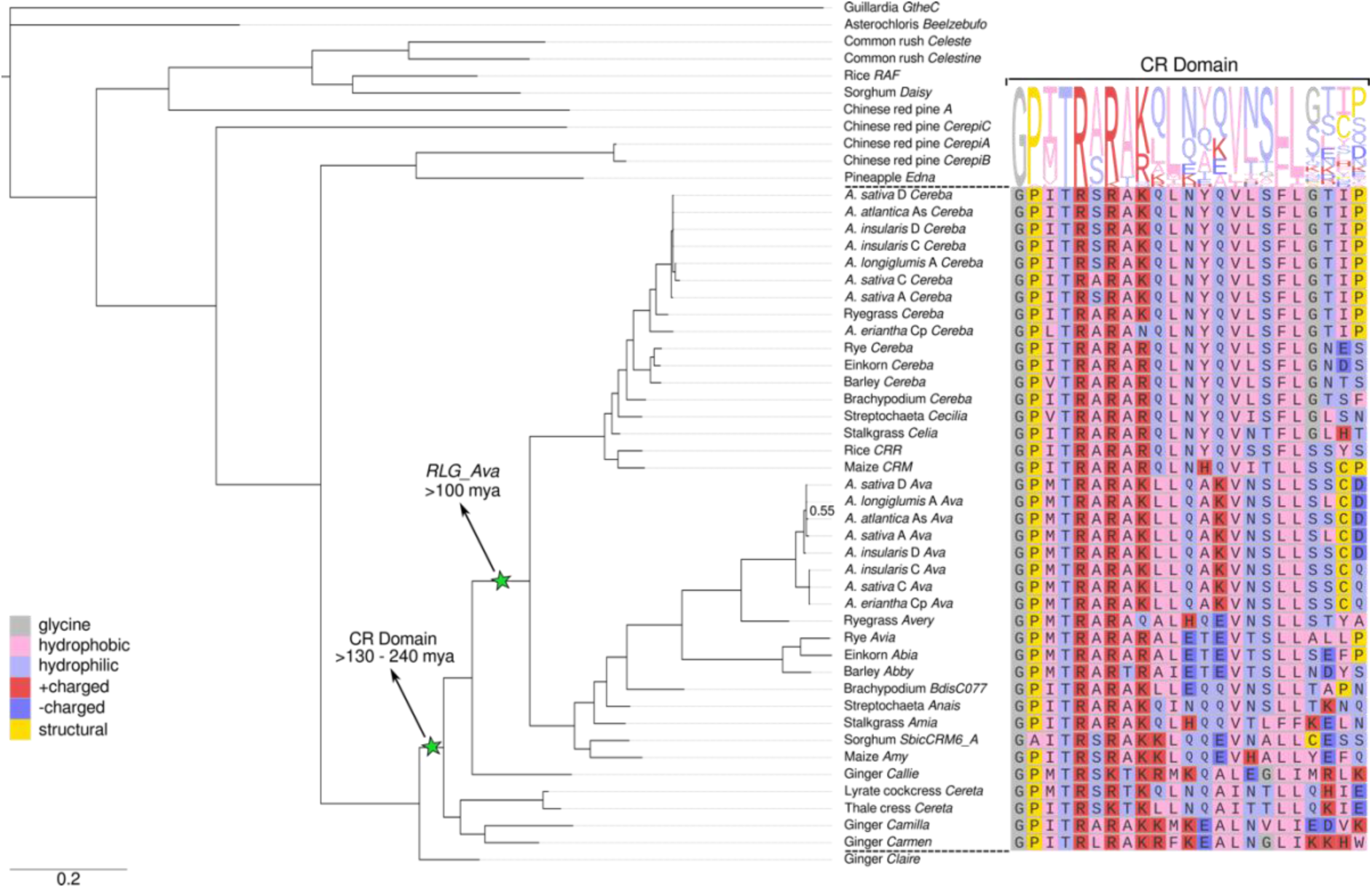
Phylogenetic tree of predicted proteins of *RLG_Cereba* and *RLG_Ava* and homologous TE families. The tree was built from the RH, RT and INT protein domains of the different TE families using MrBayes (generations= 235000, burnin of 25%). The green stars indicate the rise of the CR domain and the emergence of the *RLG_Ava* homologs in grasses (*Poaceae*). The *Guillardia theta* (GtheC) sequence was used and set as the outgroup in the MrBayes analysis. The CR-domain sequences with the conserved motive are displayed next to the respective consensus.

When focussing on the CR-domain, we identified a clear pattern of conservation of the previously described RARA[AK]-motive (here R[AS]RA[RK]), especially of the positively charged amino acid positions which strengthens the hypothesis of an important role of this sequence motive in the centromere specificity of these elements recently presented by Heuberger et al. [18] (Fig. 4). Further, we found this motive to be conserved not only across the *Poaceae* but further in ginger and interestingly also in the dicot *Arabidopsis*. In the *RLG_Claire* element from ginger we were not able to detect a CR-domain (Fig. 4). However, despite detailed analysis we can not exclude the possibility of an unresolved frameshift due to suboptimal quality of the consensus, causing this domain to not be detected. Most of the analyzed sequences displayed in Figure 4 without a CR-Domain, contain a CHROMO (<u>CHR</u>omatin <u>O</u>rganization <u>M</u>odifier) domain, which we identified using NCBI conserved domains. Merely the *RLG_CerepiC* (identified in Chinese red pine) and *RLG_Daisy* (identified in *Sorghum*) families did not exhibit this additional domain.

As we find the CR-domain present in phylogenetically distant plants, its origin must lie in the common ancestor of monocotyledons and dicotyledons more than 130 to 240 mya [53,54]. To investigate the origin of the CR-domain we built a hidden Markov model (HMM) to search against *Arabidopsis* proteins (TAIR v10.1). However, we were not able to identify a protein with a similar domain, indicating that the CR domain did not evolve from a known plant protein. Further, a search against the NCBI CD database did not yield a result.

Although there are instances in which the CR domain is missing from centromeric retrotransposons, for example in CRM (maize) group B elements [20], the conservation of the CR-domain, specifically the positively charged amino acids of the R[AS]RA[RK]-motive, for a span of over 100 myrs greatly emphasizes its importance for the centromere specificity of TEs.

## Discussion

In this study, we analyzed centromere sequence composition and organization in multiple *Avena* species. Hexaploid oat has one of the largest genomes sequenced so far, and centromeres in large-genome crops are notoriously difficult to sequence [16,21,22]. The more relevant it was that we had access to high-quality genome assemblies from several species of the *Avena* genus. On the one hand, this allowed for in-depth analysis of individual centromeres, and on the other hand, for comparative analyses of centromeres between species.

In general, we found that functional centromeres largely coincide with the insertion sites of centromere-specific retrotransposons. This mirrors previous studies in wheat [16,21], which showed that the youngest centromere-specific TE copies are mostly found inside or close to the functional centromeres. The *Avena* centromeres are mostly composed of three LTR retrotransposon families, *RLG_Cereba*, *RLG_Ava* and *RLG_Beth*. *RLG_Cereba* and *RLG_Ava* are autonomous TEs which encode all proteins necessary for their replication as well as a CR domain that presumably guides insertions to functional centromeres through targeting of centromeric CENH3 histone variants [18]. In addition, we found that *RLG_Beth* is a non-autonomous element that is likely cross-mobilized by *RLG_Ava*. This is surprisingly similar to the autonomous/non-autonomous pair *RLG_Cereba* and *RLG_Quinta*, which shape the centromeres in Einkorn wheat [18]. However, *RLG_Beth* and *RLG_Quinta* have no sequence homology and thus must have evolved independently.

Despite these similarities, we made several findings that distinguish *Avena* centromeres from each other and from previously described plant centromeres, emphasizing the importance of comparative analyses for evolutionary studies. These findings are discussed below.

### Two retrotransposon families compete for the centromeric niche in *Avena* and its relatives

Our phylogenetic analysis showed that the evolutionary *RLG_Ava* and *RLG_Cereba* lineages had been present already in the common ancestor of the grasses (*Poaceae*), which dates their origin to ∼100 mya [52]. Thus, the two retrotransposon lineages must have coexisted in grasses since then, competing for the centromeric “niche”. Interestingly, our data indicate that the dynamics can change rapidly in evolutionary terms: the *Avena* C genomes are practically void of *RLG_Cereba* elements, meaning that they had become largely silent after the C genome lineage diverged from the A/D lineage ∼8 myr [26]. This is similar to findings in the Triticeae tribe (wheat, barley and rye) where the *RLG_Ava* lineage went largely silent, as mostly fragments of *RLG_Abia* (the *RLG_Ava* homologs) were found in wheat [17]. In contrast, the *Avena* A and D genome lineages show recent activity of both *RLG_Ava* and *RLG_Cereba*, indicating that both can persist simultaneously at similar abundance in a given species. This raises the question why none of them went completely extinct in a given species or species groups and how they were able to persist over long evolutionary times. Even if one of them goes largely silent, a few functional copies must be maintained, allowing re-emergence of the TE family at a later point in time. Another possibility is that there is occasional horizontal transfer (HT) of centromeric retrotransposons between species. Indeed, multiple studies suggested that HT might be frequent, especially between closely related species [55]. This may allow TEs to circumvent silencing mechanisms (e.g. through small interfering RNAs) that were established in a given species [56]. However, we did not find any of the phylogenetic incongruencies that are defining characteristics of HT, and therefore see it more likely that the respective TE families persisted at low abundance.

### Retrotransposon populations shed light on the evolutionary history of *Avena* species

Molecular dating of TE activity bursts showed that different retrotransposon subfamilies emerged and were active at different times and in different *Avena* genomes. This informed us on the relationship between *Avena* genomes and the timing of divergences and polyploidization events. For example, the *Avena* A genomes contain multiple lineage-specific subfamilies of *RLG_Cereba* and *RLG_Ava* retrotransposons that were highly active in very recent evolutionary times, giving individual A genome species highly distinct TE profiles. This showed that the A genome of *A. sativa* is clearly distinct from both the other A genomes species *A. longiglumis* and *A. atlantica*., representing a separate evolutionary lineage. Thus, our results support the phylogeny proposed by Peng et al. [26].

Furthermore, insertion ages of retrotransposon subfamilies indicated that tetraploidization occurred at least 440,000 years ago, which is in agreement with previous findings [26]. Additionally, we date the hexaploidization event to at least 150,000 years ago. Interestingly, analysis of *RLG_Aurora* (a retrotransposon family that is not centromere-specific) indicated that there must have been a time span of ∼300,000 years between formation of the tetraploid and the hexaploid, because *RLG_Aurora* had an activity burst that is found exclusively in the tetraploid (CD) and the CD genome of hexaploid oat. This family must have been largely silent before tetraploidization, as well as at the time when the hexaploid formed. Thus, *RLG_Aurora* might be a rare remnant of a “genomic shock”, a dramatic activation of TEs, that is postulated to follow polyploidization [57,58]. However, no evidence for genomic shock was found in other polyploids such as wheat [17], nor did we find any similar TE activity burst following hexaploidization.

Here, we want to emphasize that for all our molecular dating procedures, we used the nucleotide substitution rate of 1.3E-8 per site per year, which is typically used to date insertion ages of retrotransposons [59]. Our estimates for species divergences and polyploidization events tend to be somewhat more recent than those done with other methods (e.g. [26,60]). Nevertheless, our results are overall consistent with those of previous studies.

### *Avena* centromeres show evolution of different centromeric architectures

One of the most curious findings of our study were the strikingly different centromere architectures in the A, C and D genome lineages. It has long been known that plants have a variety of centromere architectures [61]. In *Arabidopsis*, for example, centromeres largely consist of short tandem repeats that are occasionally interspersed with TEs [62], while in maize and wheat, they are largely comprised of CRM/Cereba-type retrotransposons [11,16]. Intermediate forms that contain both TEs and tandem repeats in abundance are, for example, found in Brachypodium [63]. Nevertheless, we were surprised to find such variety in a genus that diverged probably less than∼ 8 myrs ago [26].

While the centromes of the A and D genomes are nearly exclusively comprised of LTR retrotransposons of the *RLG_Cereba*, *RLG_Ava* and *RLG_Beth* families, those in the C genomes lack *RLG_Cereba*, but contain a high number of tandem repeats. Furthermore, the three lineages show very different TE activity profiles: the A genomes contain multiple TE subfamilies that were very recently active, and may, in fact, still be active now. In contrast, the D genomes contain multiple subfamilies that were highly active within the past million years, but seem to be silent currently. Furthermore, only one of the identified subfamilies (*RLG_Cereba_G1_2*) was found in both the A and D genomes. This indicates that centromere compositions can vary strongly even in very closely related species.

However, the most dramatic differences were found in the *Avena* C genomes. Not only do *RLG_Cereba* elements seem to have gone nearly extinct, the *RLG_Ava* retrotransposons have been largely silent for at least the past million years. Additionally, we found that ∼17% of centromeric sequences are composed of tandem repeats of a short 46-51 bp motif. While a previous study reported short tandem repeats of that length to be centromeric [25], we find this sequence motif in hundreds of thousands of copies across the entire C genome. This highly repetitive sequence must have emerged in the C genome ancestor, as it is found in high abundance in *A. sativa*, *A. insularis* and *A. eriantha*. We have no hint as to the origin of this sequence since it has no homology to known TE sequences. Neither do we know whether the emergence and spread of these tandem repeats have any causal connection to the silencing of the centromeric *RLG_Ava* retrotransposons. However, our data indicate that CENH3 deposition in the C genome is less strictly associated with insertion sites of the youngest centromeric retrotransposons than in the A an D genomes. This could suggest that other (e.g. epigenetic) factors influence positioning of centromeres in the C genome. We acknowledge that we projected the position of the centromeres in *A. sativa* OT3098 from positions published for *A. sativa* cv. Sang [42]. This strategy can merely provide estimations, as recently demonstrated for the *T. aestivum* accessions Chinese Spring and Julius [21].

### On the origin of centromere-specific retrotransposons in grasses

All centromere-specific retrotransposons in grasses described so far are of the CRM/Cereba-type which have in common a distinct CR domain fused to their integrase. Recent studies indicated that the CR domain may indeed interact directly with centromeric CENH3 histone variants and thus guide new insertion to the functional centromere [18]. It has long been known that TEs sometimes acquire protein domains from cellular genes, which gives them new functions. For example, a number of retrotransposons were shown to have acquired chromodomains which are known to recognise histones [20]. However, the origin of the CR domain is still unclear as it lacks homology to canonical chromodomains [13]. In this study, we searched plant protein databases with HMMs of the CR domain but did not find any homologs. One explanation is that the CR domain is so highly diverged that its origins are not detectable anymore. Alternatively, the CR domain could have been acquired through horizontal transfer from a different organism group (e.g. bacteria). However, our search of conserved domains at NCBI also did not yield any results. It is still possible that the CR domain was acquired from an unknown source organism that is not represented in the conserved domain libraries, as previously suggested [13]. As a third possibility, we hypothesize that the CR-domain may represent a true evolutionary innovation. It could have been gained by chance through an extension of the canonical reading frame (i.e. through loss of a stop codon) in the *RLG_Ava* and *RLG_Cereba* ancestral TE, aligning with the general idea of a spontaneous acquisition recently proposed by Neumann and colleagues [13]. At this point we favour this last hypothesis.

## Conclusions

The high-quality assemblies of multiple genomes from the *Avena* genus allowed for detailed comparative analyses of centromeres and their evolutionary dynamics in these highly repetitive genomes. We showed that *Avena* centromeres are largely composed of two retrotransposon families, but also found striking differences in centromere architecture and sequence composition. These included the emergence and disappearance of retrotransposon families and subfamilies, centromere shifts and the spread of novel satellite repeats. Considering that the different species studied here all belong to the same genus and are closely related, our data show that centromeres are highly dynamic and can rapidly take different evolutionary trajectories. Thus, continuing to produce high quality assemblies especially from large and complex crop genomes will provide valuable resources to investigate the molecular mechanisms that drive centromere evolution on a more broader evolutionary scale.

## Methods

### TE annotation

The TE annotation was performed by running EDTA (v2.2.1 [49]) on the OT3098 v2 genome using the settings --overwrite 1 --sensitive 1 --anno 1 --force 1. The TREP database v.19 (Schlagenhauf and Wicker, 2016 available at https://trep-db.uzh.ch) was used as a curated library input and further was utilized to manually annotate the identified high copy sequences on a family-level by blasting the sequences against the database. In addition, the EDTA pipeline (v2.2.1 [49]) was run on the estimated centromere region segments from the OT3098 v2 genome with the settings mentioned above. We provided a curated library combining specifically Poaceae TEs filtered from the TREP database (Schlagenhauf and Wicker, 2016 available at https://trep-db.uzh.ch) and grass TEs which were manually curated during the course of this study.

### TE population analysis

Following the pipeline described in Wicker et al. [45], full length elements were identified starting with the isolation of around 100 LTRs per family. Next the LTRs were aligned using Clustalw with a gap opening penalty of 10 a gap extension penalty of 0.2. If variants of LTRs were visible as groups in the alignment, group specific consensus sequences were constructed. These consensus LTRs were then blasted against the chromosomes of *A. sativa*. If two LTRs were found in the same orientation, within a range of ±1000 by of the original consensus and not more than 5 bp were missing from either of the LTRs (checking for completeness), an identified element was classified as full length. An additional filtering step was applied to remove elements with large insertions from deletions. In addition, blast searches against the TREP database (Schlagenhauf & Wicker, 2016 https://trep-db.uzh.ch) were performed to confirm the family identity. The full length TEs were then used in the following analysis. To estimate the insertion age of the individual TEs we used an in-house script. It aligns the LTRs using the program Water from the EMBOSS package, distinguishes between transitions and transversions and applies a rate of 1.3E-8 per site and year for the molecular dating [44,59]. For the visualization the age of elements with estimated age older than the 95% quantile of the respective family or displayed subset of a family were set to 95% quantile value to ensure good visible resolution of the age distribution. The individual full-length copies were aligned against the initial consensus using the program Water (EMBOSS package) with a gap opening penalty of 50 and a gap extension penalty of 0.1. We used an in-house script to combine the pairwise comparisons into one variant call format (vcf) file. Variants with an occurrence of less than 5% were filtered out (minor allele frequency set to 5%). To calculate the PCA we used the R packages SNPRelate (v1.38.0, [64]) and gdsfmt (v1.40.0, [64]).

To display the PCA for identification of the TE subfamilies an in-house perl-script was used. The subfamilies were defined by visual inspection of the plot within the user interface following the TEpop pipeline. The 95^th^ percentile of the estimated insertion age per identified subgroup was used as a cut off for copies to include in further analysis of these groups as estimated insertion ages above this threshold are likely artefacts. For example, they can be the result of a crossing over of two TEs thus leading to pairing of two unrelated LTR sequences. In the following, these insertion ages were used for the divergence time estimation. For violin plots only groups with less than 10 entries were not included. For the visualization R (R Core Team 2024) and the packages stringr (v1.5.1 https://CRAN.R-project.org/package=stringr), ggplot2 (v3.5.1 [65]), dplyr (v1.1.4 https://CRAN.R-project.org/package=dplyr) and tidyverse (v2.0.0 [65]).

### Centromere position estimation

Centromere positions were predicted using an in-house perl script to compare segments of *A. sativa* cv. Sang and A. sativa OT3098 genomes. To estimate the region for comparison the position of centromeres from *A. sativa* cv. Sang were taken from the work of Liu and colleagues [42]. A sequence of around 100Mb per chromosome from the Sang genome surrounding the published positions was compared using a blastn search to the corresponding full chromosome of OT3098 by blasting 1000bp segments with a sliding step of 10000. Centromere positions on OT3098 were then projected from the positions which were matched with the centromere position in Sang. In addition, visual comparison of the alignment of the defined segment of the *A. sativa* cv. Sang genome and the positions on *A. sativa* OT3098 genomes were performed.

### Phylogenetic analysis of centromere specific retrotransposons

First we used Clustal W [66] to align the consensus sequences and created a cropped alignment using an in-house script. For the following analysis we used a subset of the full length polyproteins including the RT, RH and INT protein domains, whose position was estimated beforehand using NCBI conserved domain batch search with standard settings. We used Clustal X [66] to transfer the alignment into nexus format and ran MrBayes [51] 25000 generations with a burnin value of 25%. The data visualization was done using R (R Core Team 2024) and the packages stringr (v1.5.1 https://CRAN.R-project.org/package=stringr), ggplot2 (v3.5.1 [65]), ggmsa (v1.3.4 [67,68]), ape (v5.8 [69]), ggtree (v3.10.1 [68]), ggtreeExtra (v1.14.0, [70]).

### Hidden Markov models (HMM) for identification of CR-domain origin

To search for proteins similar to the CR-domain we build a HMM from an alignment of the sequences which were also used to build the phylogenetic tree, cropped to the CR-domain. Using HMMER (v3.4 (2023) available under http://hmmer.org/) we built the model with the function hmmbuild and searched the proteins of *A. thaliana* Tair v10.1 using hmmsearch.

### Tandem-repeat analysis

To identify tandem repeats in centromeres, we defined the centromeres by using a running average of RLG_Ava insertion ages. The local minimum was taken as centromere midpoint, to which 5 Mb on each side were added. The putative centromeres were then annotated with tandem repeats finder (trf, Benson et al.) using the settings: Match=2, Mismatch=7, Delta=7 PM=80, PI=10, Minscore=50, MaxPeriod=2000.

## Abbreviations

AP: Aspartate protease
CDS: Coding sequence
ChiP: Chromatin Immunoprecipitation
CR: Chromodomain
FISH: Fluorescence in-situ Hybridization Assays
INT: Integrase
LTR: Long Terminal Repeats mya Million years ago
myrs: Million years
PCA: Principal component analysis
RH: RNase H
RLG: Retrotransposon LTR Gypsy
RT: Reverse transcriptase
TE: Transposable Element

## Declarations

### Ethics approval and consent to participate

Not applicable

### Consent for publication

Not applicable

### Availability of data and materials

The datasets analyzed during this current study are available in the GrainGenes repository (*A. sativa* cv. Sang v1.1 https://wheat.pw.usda.gov/GG3/content/avena-sang-download;

*A. sativa* OT3098 v2, PepsiCo, https://wheat.pw.usda.gov/jb?data=/ggds/oat-ot3098v2-pepsico) and the NCBI Genome Database (*A. longiglumis* CN58138 v1 <u>GCA_910589755.1</u>*, A. eriantha* BYU132 v1 <u>GCA_910589775.1</u>, *A. insularis* BYU209 v1 <u>GCA_910574615.1</u>, *A. atlantica* CC7277 v1, *Arabidopsis thaliana* TAIR v10.1 <u>GCF_000001735.4</u>). The produced TE consensus sequences were deposited in the TREP database (Schlagenhauf & Wicker, 2016 https://trep-db.uzh.ch).

## Authors’ contributions

CMW analyzed the centromere specific transposable elements with input from MH and TW. MH performed tandem repeat analysis with input from TW. TW. designed the research. CMW and TW wrote the manuscript with input from MH.

## Acknowledgements

We would like to thank S3IT and the IT-Team of IPMB for their continued support.

