## supplementary_material for "Comparative analysis of centromeres of oat (*Avena sativa*) and its tetraploid and diploid relatives reveals rapid evolution of centromere composition and architecture"

### Supplement

#### Supplementary Figures

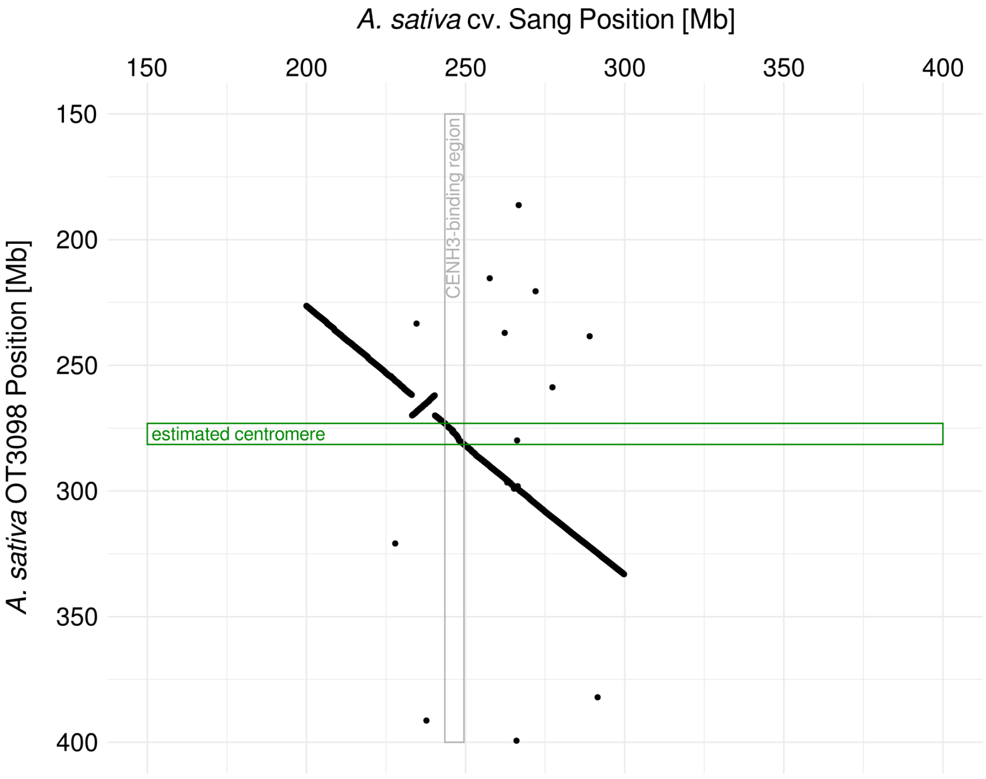


**Figure S1.** Results of blast comparison of the pericentromeric regions of chromosome 1C from *A. sativa* cv. Sang and *A. sativa* OT3098 for centromere position estimations.

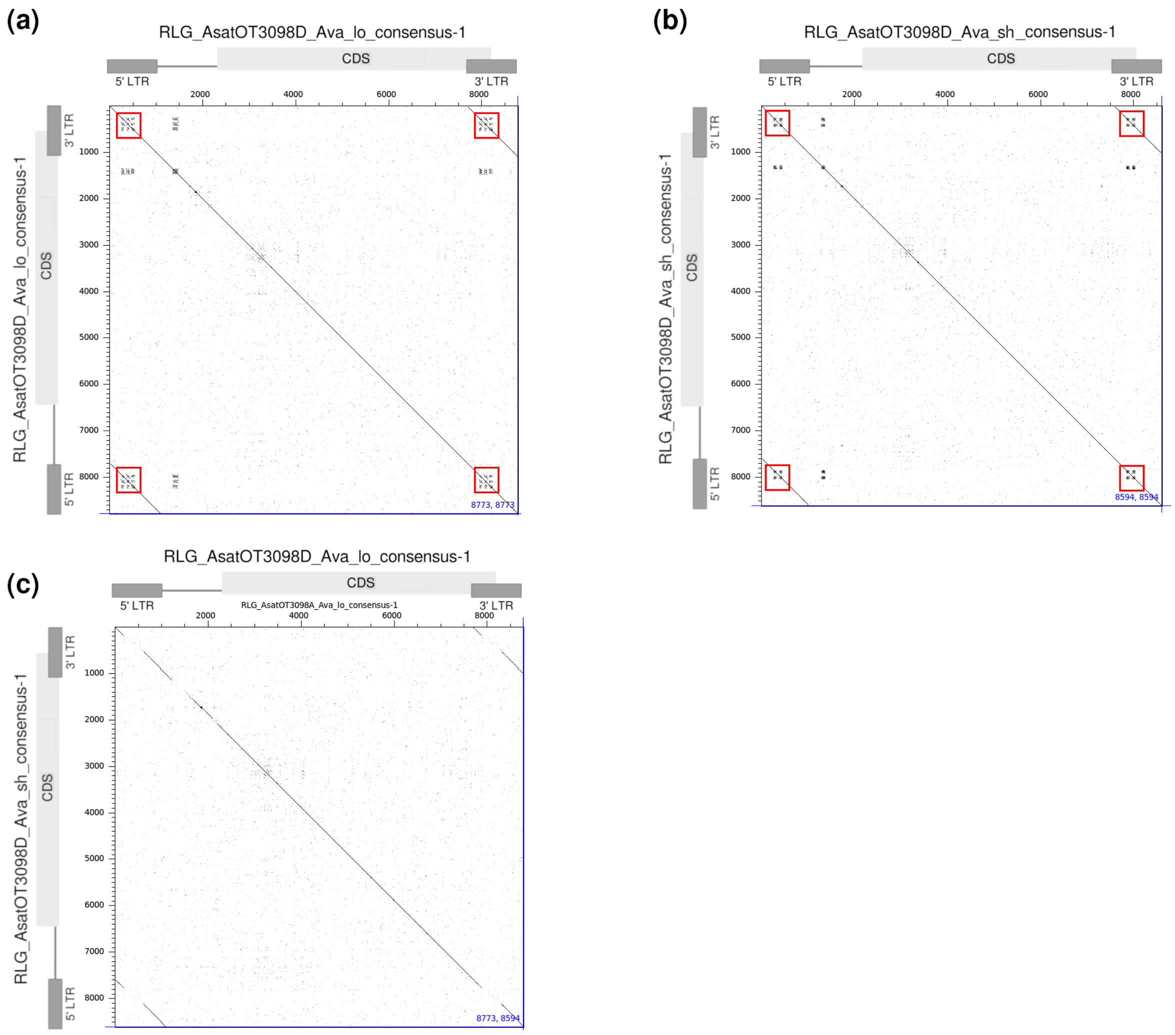


**Figure S2.** Dotplot comparisons of variants of RLG_Ava retrotransposons. (a) Dot plot alignment of the longer variant RLG_AsatOT3098D_Ava_lo_consensus-1 against itself. *RLG_Ava* elements contain small tandem repeat clusters in their LTRs (indicated by red boxes) which differ among the main variants. (b) Dot plot alignment of the shorter variant RLG_AsatOT3098D_Ava_sh_consensus-1 against itself. (c) Dot plot alignment of the two variants. Note that the tandem repeat arrays show no DNA sequence conservation while the rest of the sequence is well conserved.

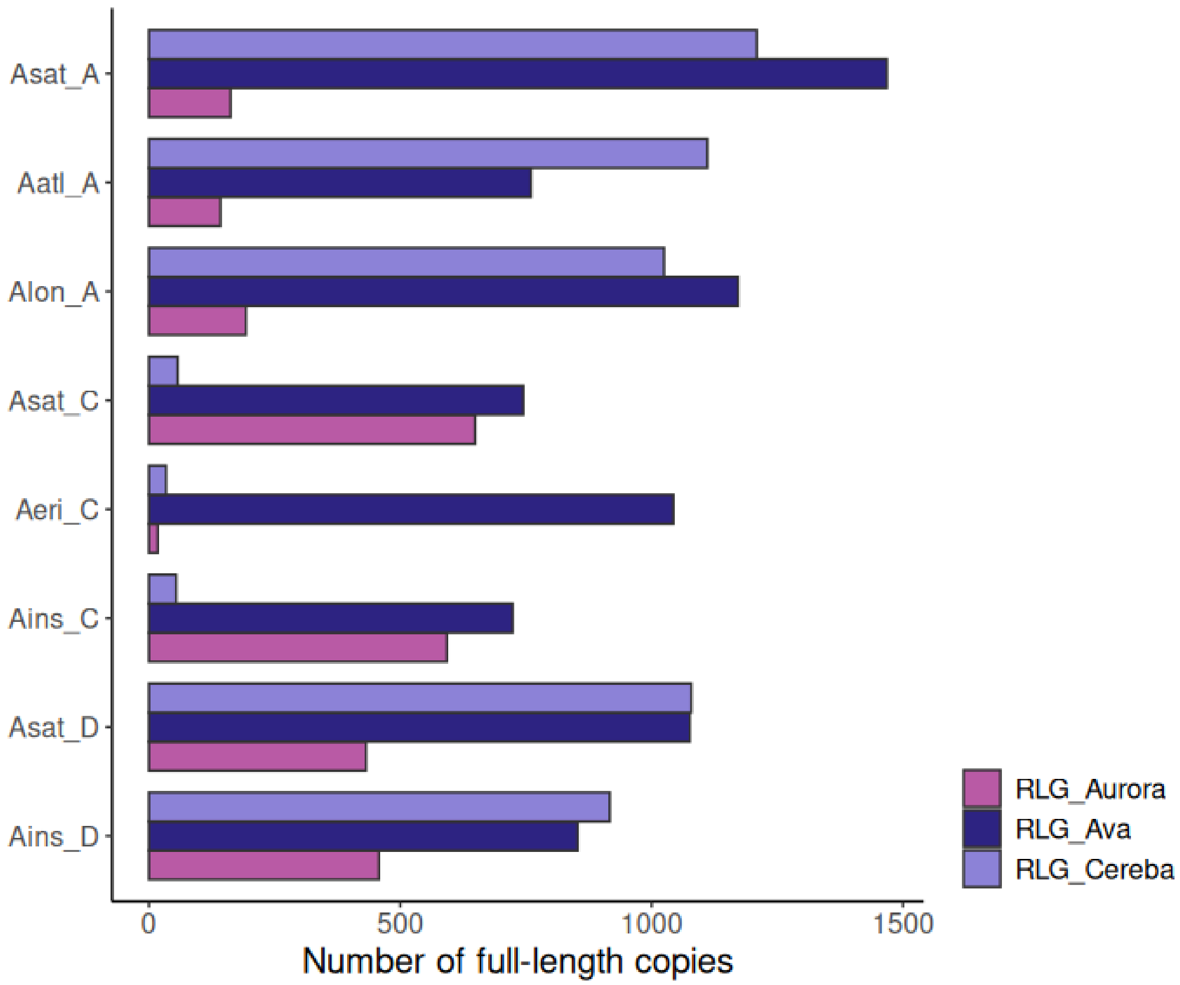


**Figure S3.** Number of identified full-length copies of *RLG_Ava*, *RLG_Cereba* and *RLG_Aurora* in different genomes.

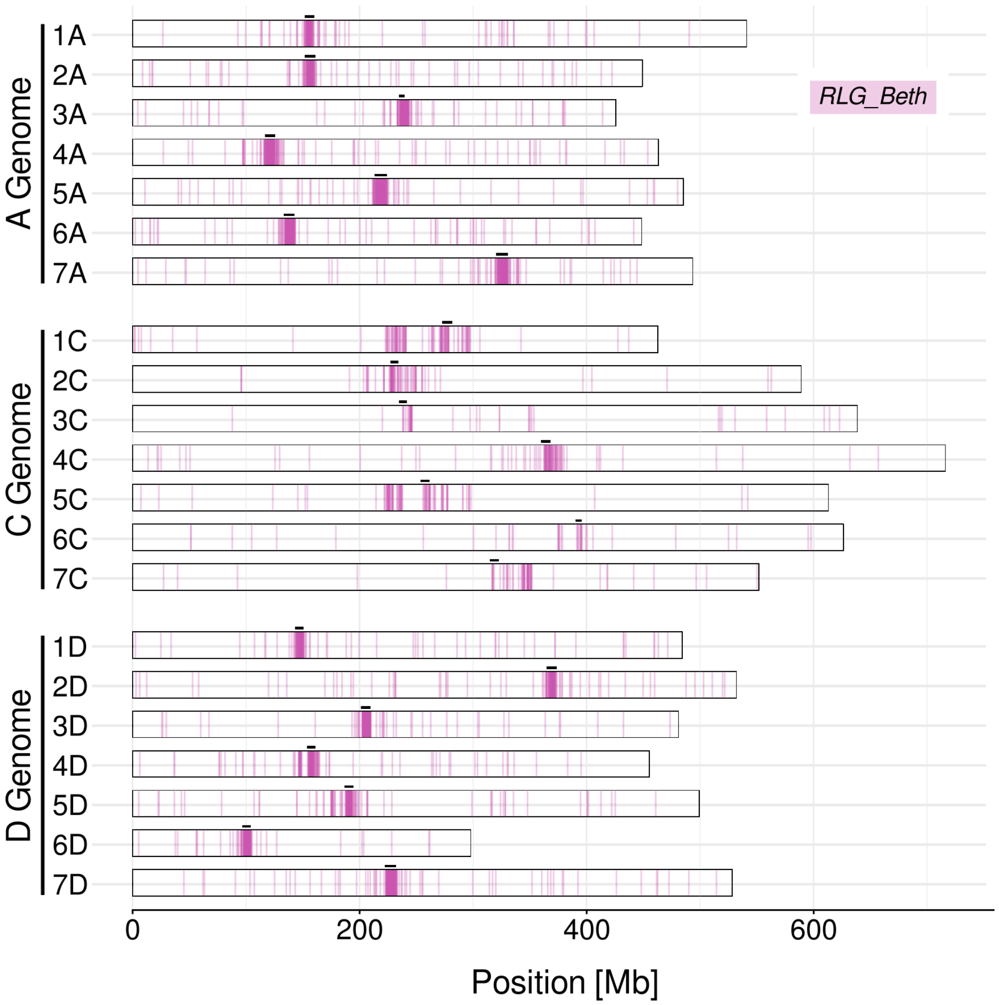


**Figure S4.** Localization of full-length copies of *RLG_Beth* retrotransposons across the *A. sativa* OT3098 genome. Positions of functional centromeres inferred from CENH3 ChIP-seq data are indicated by horizontal black bars.

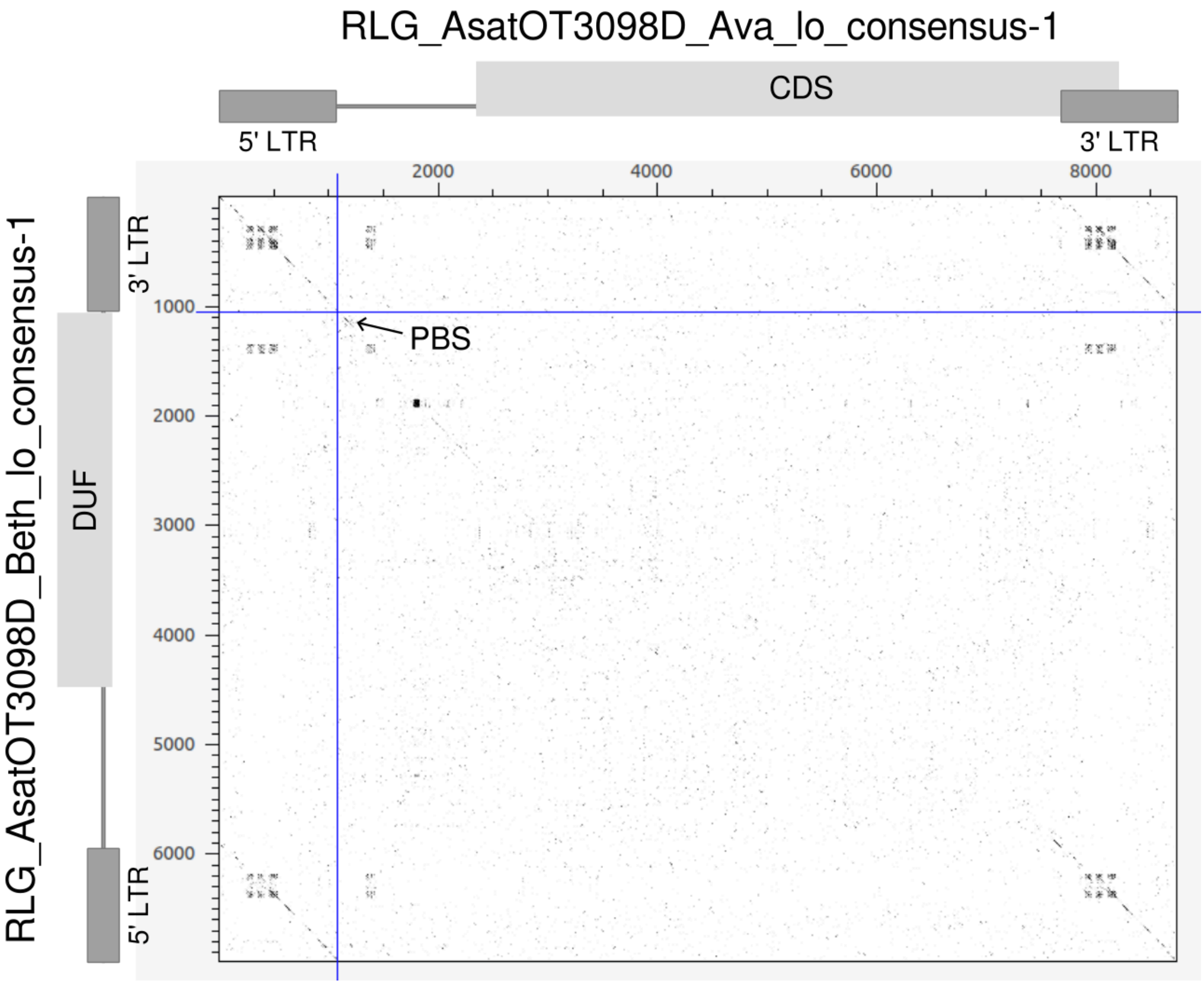


**Figure S5.** Dotplot comparison of *RLG_AsatOT3098D_Ava_lo_consensus-1* and *RLG_AsatOT3098D_Beth_lo_consensus-1*. *RLG_Ava* encodes the typical retrotransposon proteins necessary for replication, such as reverse transcriptase, integrase etc. while *RLG_Beth* only encodes a domain of unknown function (DUF). LTRs and primer binding site (PBS) are conserved between the two, suggesting that the autonomous *RLG_Ava* retrotranspsons cross-mobilize the non-autonomous *RLG_Beth* elements

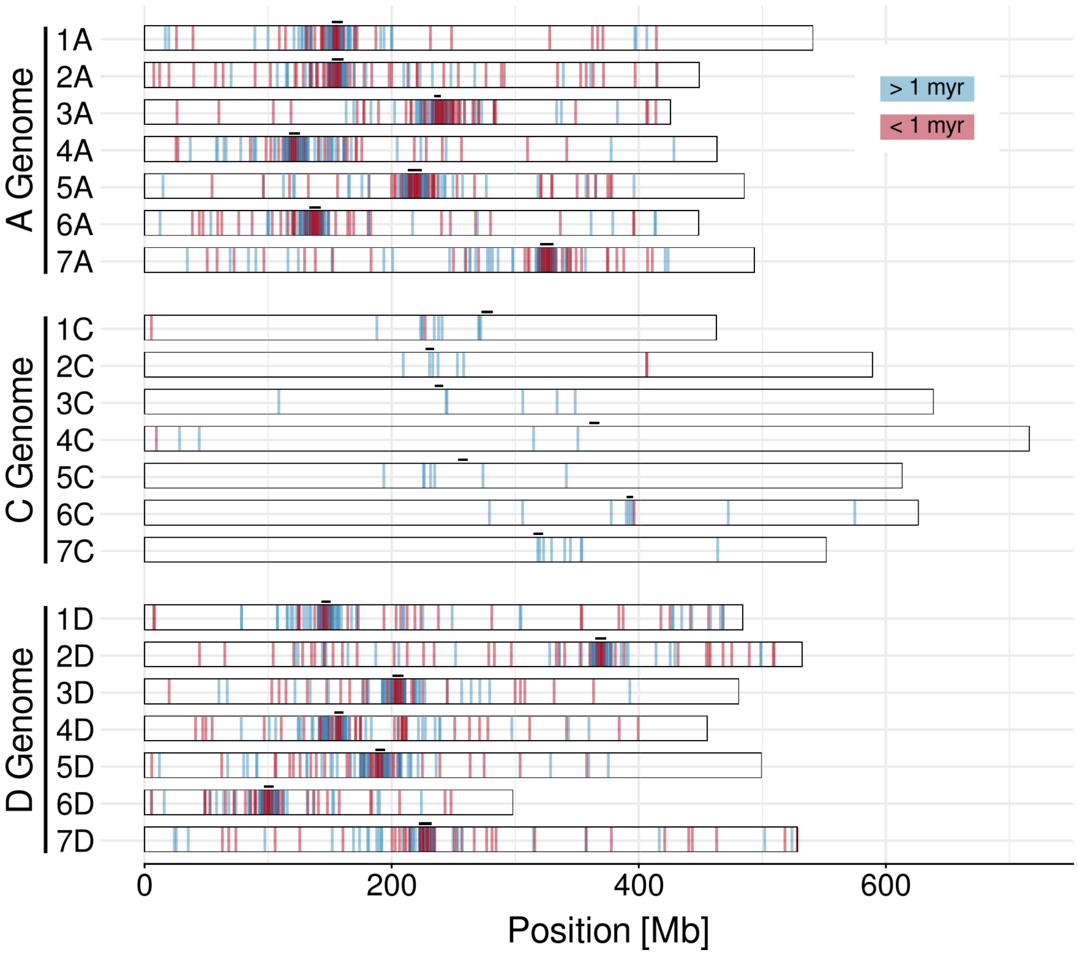


**Figure S6.** Full-length *RLG_Cereba* elements in *A. sativa* colored by age groups, young elements within insertion ages under 1 myr are displayed in red and older elements with insertion ages over 1 myr are displayed in blue. Black bars indicate the projected centromere positions.

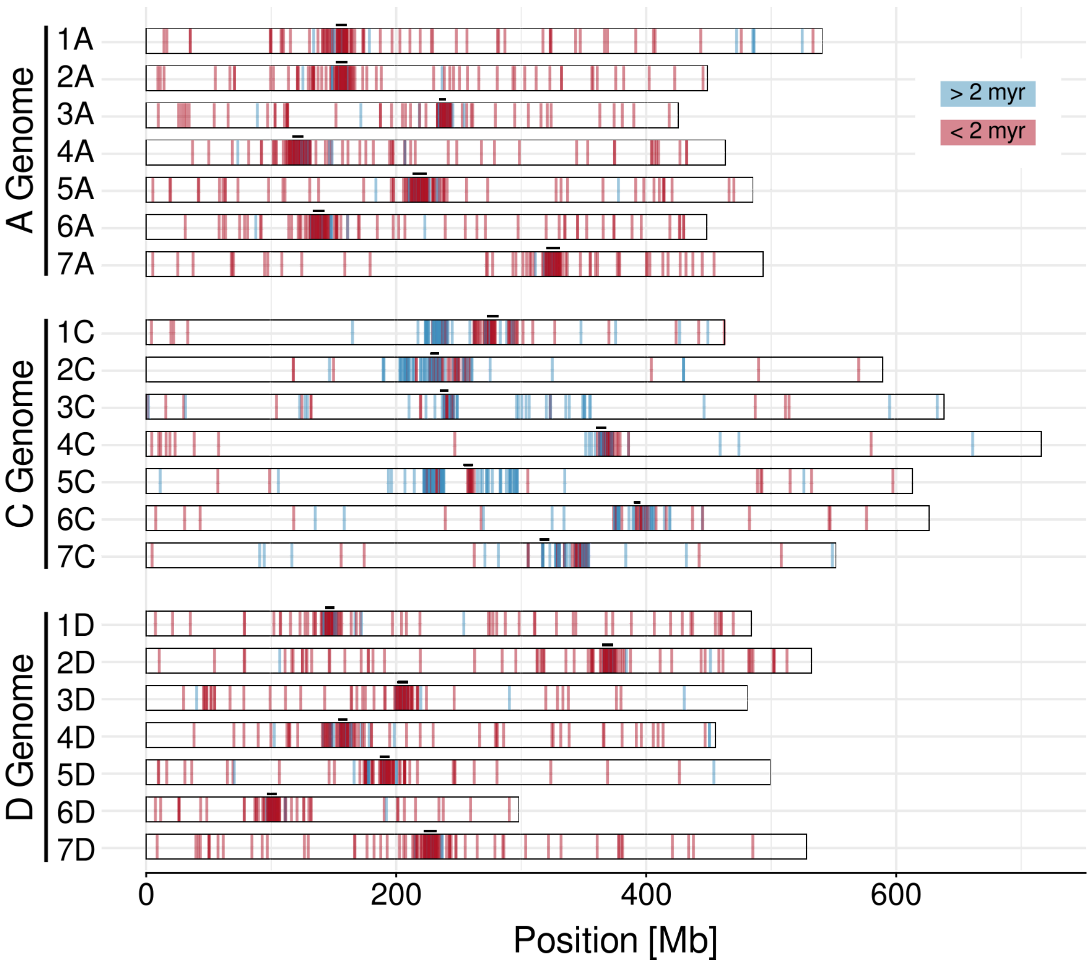


**Figure S7.** Full-length *RLG_Ava* elements in *A. sativa* colored by age groups, young elements within insertion ages under 2 myr are displayed in red and older elements with insertion ages over 2 myr are displayed in blue. Black bars indicate the projected centromere positions.

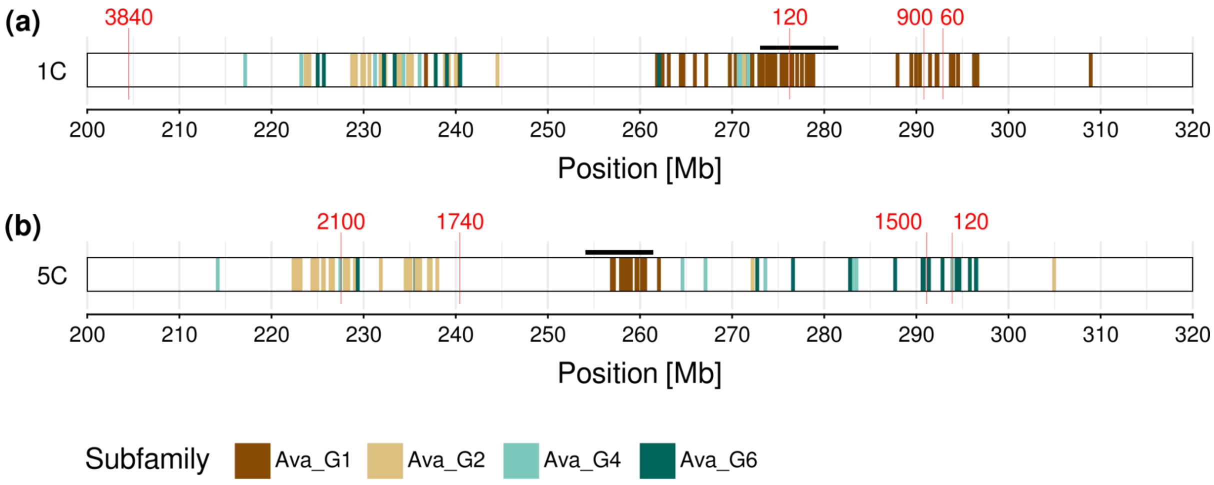


**Figure S8.** A visualization of the pericentromeric region of chromosome 1C (a) and 5C (b) with four identified subfamilies of *RLG_Ava*. Assembly gaps are marked in red and labeled with the respective length.

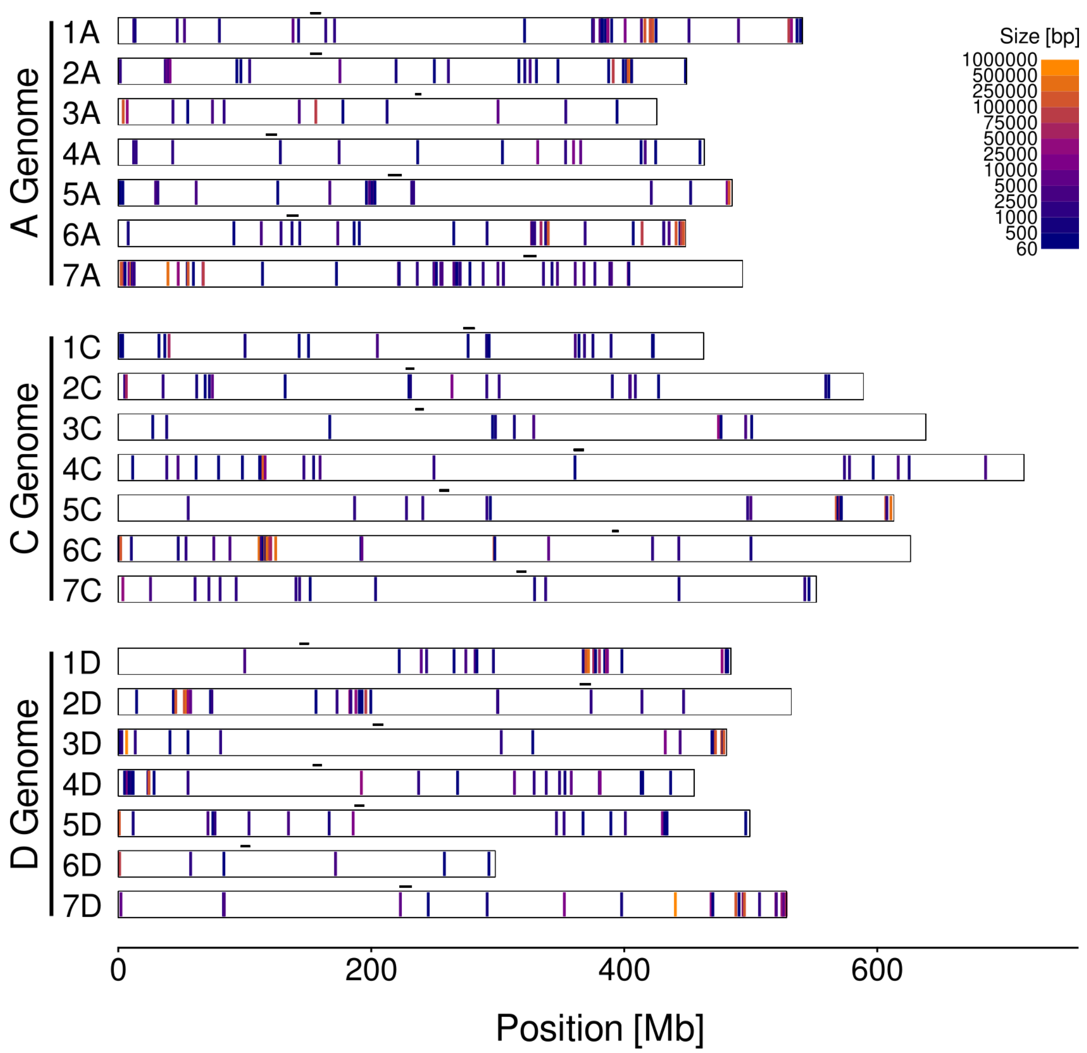


**Figure S9.** Sequence assembly gaps and estimated functional centromere position in *A. sativa* OT3098

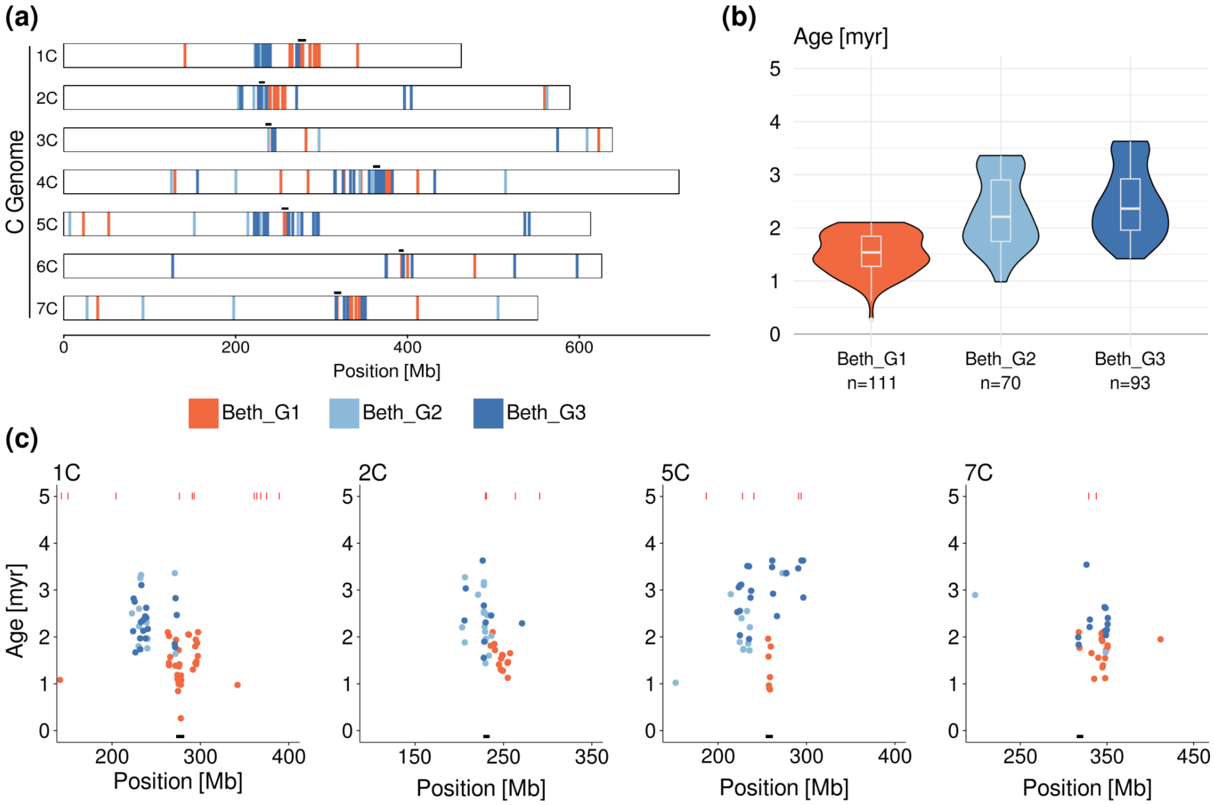


**Figure S10.** *RLG_Beth* subfamilies mark centromere shifts in *A. sativa* C genome. The distribution of copies of the non-autonomous family *RLG_Beth* (a) and the respective age distribution of the different subfamilies (b). (c) The age of *RLG_Beth* copies across the position within the centromeric region of *A. sativa*. Red marks indicate annotated gaps. The black bar indicates the estimated centromere position.

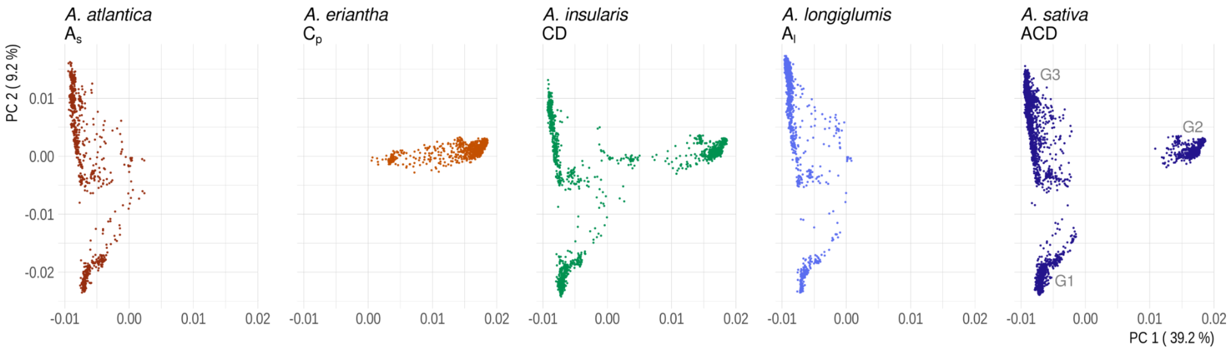


**Figure S11.** PCA of *RLG_Ava* copies by species. The three main groups are marked illustratively on the A. sativa genome PCA.

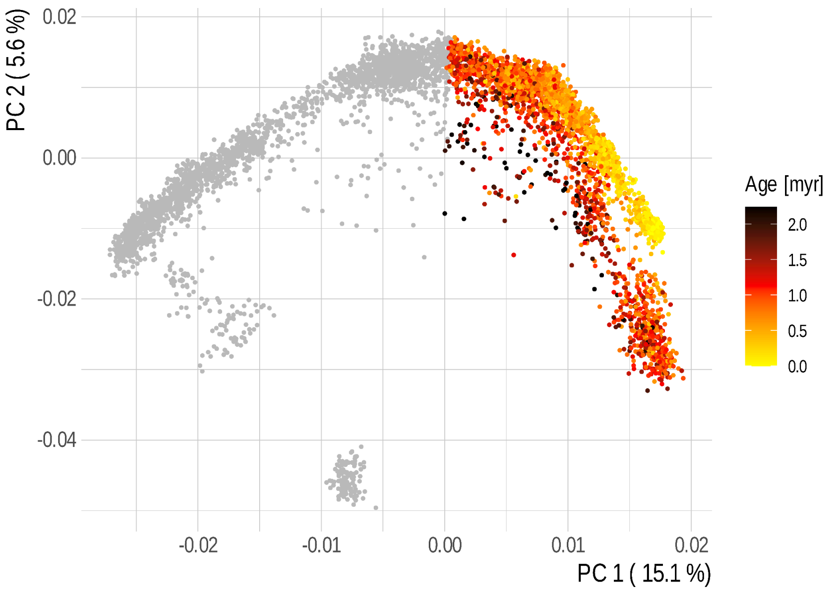


**Figure S12.** PCA of all extracted full length *RLG_Cereba* copies. The elements used for further analysis are colored in their respective insertion age.

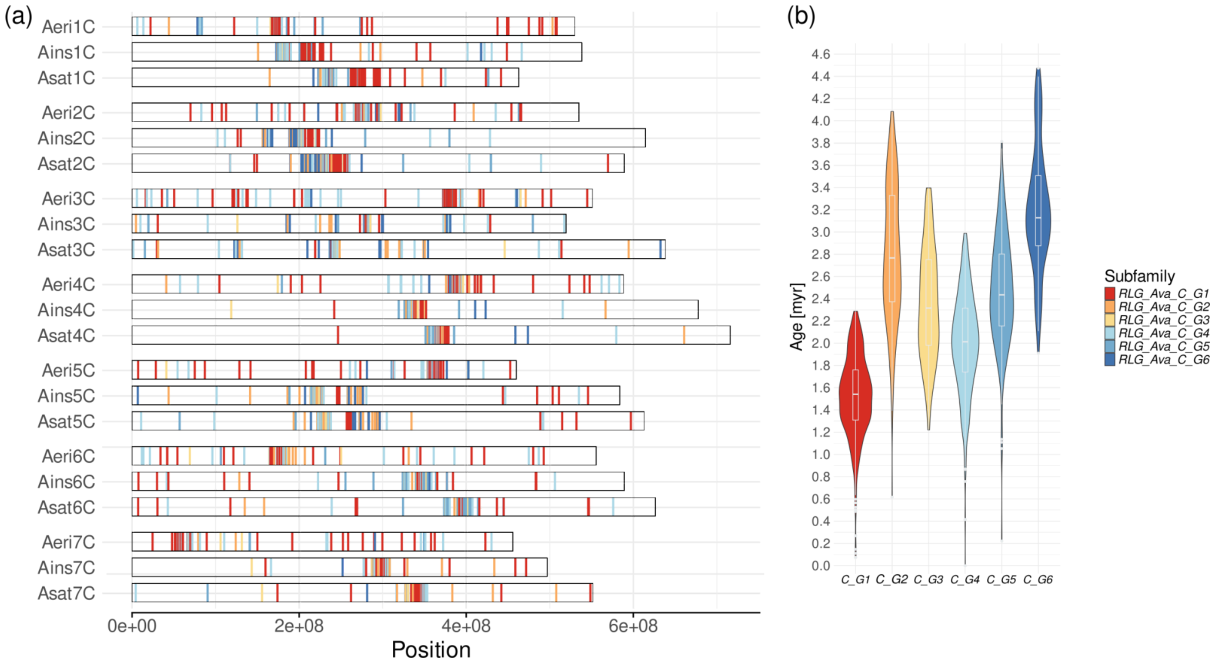


**Figure S13.** Distribution of *RLG_Ava* subfamilies on the C genomes. The *RLG_Ava* copies found on the C genomes of *A. eriantha*, *A. sativa* and *A. insularis* were used for a detailed analysis of subfamilies. We identified six subfamilies, *RLG_Ava_C_G1* through *G6*. The names of the subfamilies are shortened on the x axis of the violin plot for clarity. The centromere shifts we identified in the *A. sativa* C (sub-)genome are also found on the corresponding chromosomes in *A. insularis*.

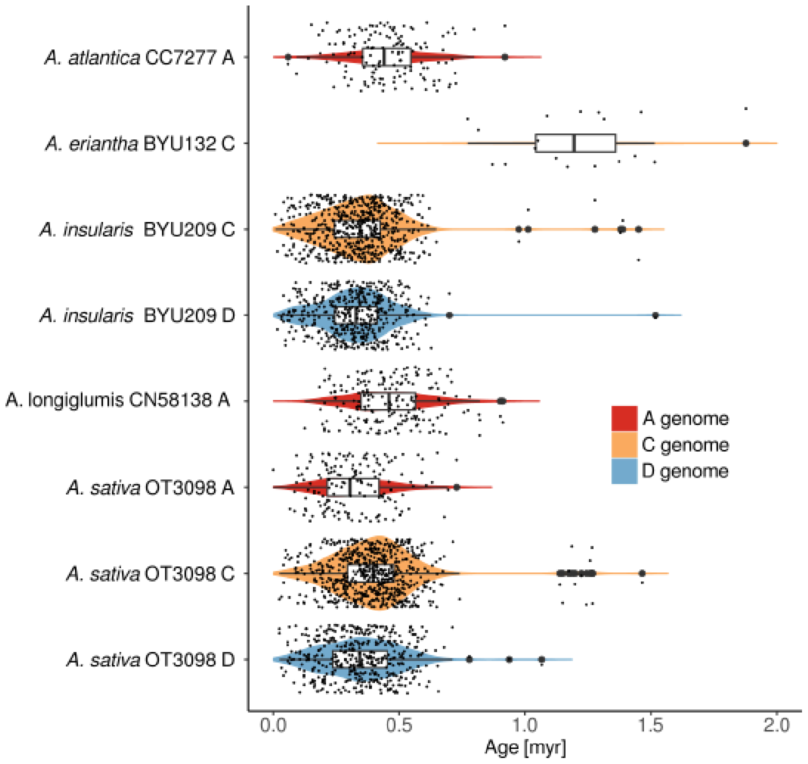


**Figure S14.** *RLG Aurora* copies across the genomes of *A. atlantica*, *A. eriantha*, *A. insularis*, *A. longiglumis* and *A. sativa*. Copies with estimated insertion ages under 2 myrs are displayed.

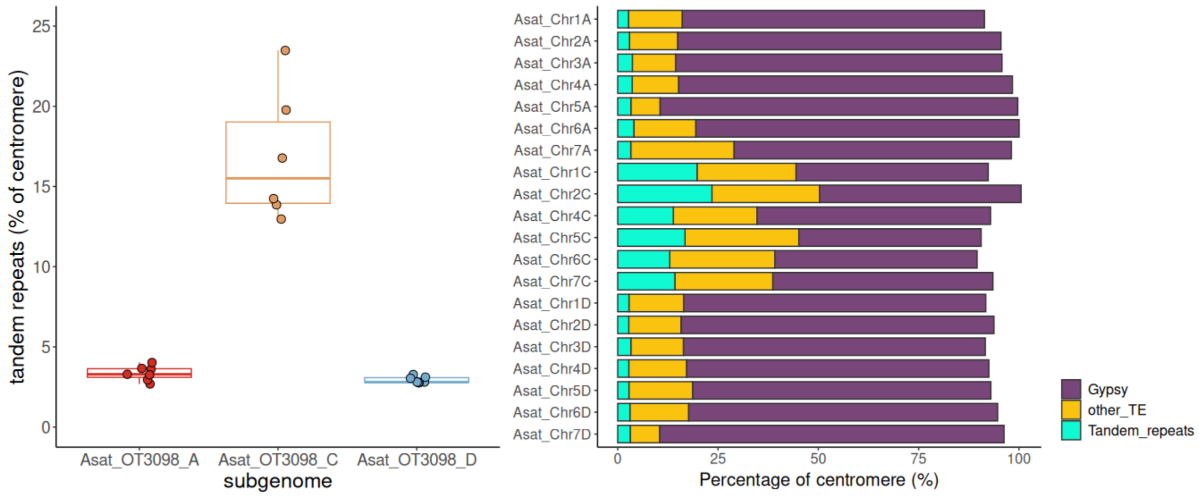


**Figure S15.** centromere composition analysis. (a) tandem repeat content of centromeres of different subgenomes of the *Avena sativa* genome OT3098. (b) composition of individual centromeres.

**Supplementary Tables**

**Table S1.** Predicted centromere positions OT3098 by comparison with *A. sativa* cv. Sang. Positions given in gray indicate mismatch between estimated centromere and main insertion site of young centromere specific elements on chromosomes.

| Chr | start | end | Chr | start | end | Chr | start | end |
|---|---|---|---|---|---|---|---|---|
| chr1A | 152.2 | 159.9 | chr1C | 273.1 | 281.5 | chr1D | 143.5 | 150.4 |
| chr2A | 152.1 | 160.6 | chr2C | 227.6 | 233.9 | chr2D | 365.1 | 373.2 |
| chr3A | 234.8 | 239.3 | chr3C | 235.1 | 241.4 | chr3D | 201.3 | 209.1 |
| chr4A | 117.1 | 125.2 | chr4C | 360.3 | 367.8 | chr4D | 154.2 | 160.6 |
| chr5A | 213.3 | 223.8 | chr5C | 254.1 | 261.4 | chr5D | 187.1 | 194.2 |
| chr6A | 133.5 | 142.2 | chr6C | 390.6 | 395.2 | chr6D | 97.0 | 104.0 |
| chr7A | 320.5 | 330.5 | chr7C | 315.2 | 322.3 | chr7D | 222.6 | 231.8 |
